# Variant-to-function analysis of the childhood obesity chr12q13 locus implicates rs7132908 as a causal variant within the 3’ UTR of *FAIM2*

**DOI:** 10.1101/2023.08.21.553157

**Authors:** Sheridan H. Littleton, Khanh B. Trang, Christina M. Volpe, Kieona Cook, Nicole DeBruyne, Jean Ann Maguire, Mary Ann Weidekamp, Keith Boehm, Alessandra Chesi, James A. Pippin, Stewart A. Anderson, Andrew D. Wells, Matthew C. Pahl, Struan F.A. Grant

## Abstract

The ch12q13 obesity locus is among the most significant childhood obesity loci identified in genome-wide association studies. This locus resides in a non-coding region within *FAIM2*; thus, the underlying causal variant(s) presumably influence disease susceptibility via an influence on *cis*-regulation within the genomic region. We implicated rs7132908 as a putative causal variant at this locus leveraging a combination of our inhouse 3D genomic data, public domain datasets, and several computational approaches. Using a luciferase reporter assay in human primary astrocytes, we observed allele-specific *cis*-regulatory activity of the immediate region harboring rs7132908. Motivated by this finding, we went on to generate isogenic human embryonic stem cell lines homozygous for either rs7132908 allele with CRISPR-Cas9 homology-directed repair to assess changes in gene expression due to genotype and chromatin accessibility throughout a differentiation to hypothalamic neurons, a key cell type known to regulate feeding behavior. We observed that the rs7132908 obesity risk allele influenced the expression of *FAIM2* along with other genes, decreased the proportion of neurons produced during differentiation, up-regulated cell death gene sets, and conversely down-regulated neuron differentiation gene sets. We have therefore functionally validated rs7132908 as a causal obesity variant which temporally regulates nearby effector genes at the ch12q13 locus and influences neurodevelopment and survival.

## INTRODUCTION

Childhood obesity affects approximately 14.7 million individuals aged 2-19 years in the United States, corresponding to approximately one in five children and adolescents^1^. The global prevalence of childhood obesity has increased substantially, rising from less than 1% to more than 7% in recent decades^2^. Obesity in childhood leads to a higher likelihood of obesity in adulthood^3^ and increases the risk of developing leading causes of poor health and early death via hypertension, metabolic disorders, cardiovascular disease, and common cancers^4^.

Monogenic cases of obesity arise from highly deleterious chromosomal deletions or mutations in crucial genes, while in contrast, common cases of polygenic obesity are driven by a combination of multiple environmental and genetic factors^5^. This genetic component to polygenic obesity can explain a large portion of obesity risk, with heritability estimates ranging from 40-85%^6^, but remains incompletely understood. However, it is known that neuronal pathways in the hypothalamus control food intake and are key regulators for both monogenic and polygenic obesity^5^. Several human stem cell-derived hypothalamic neuron models have been developed^7–11^ and used to investigate the molecular basis of body weight regulation^7,11–19^.

Genome-wide association studies (GWAS) have identified genomic regions that harbor susceptibility variants conferring common adult obesity^20,21^ and childhood obesity^22–25^ risk. An ongoing challenge in the field is to translate such GWAS loci into meaningful discoveries that can expand our knowledge of the genomic basis of complex traits. Most variants identified by GWAS reside within non-coding regions, so their underlying molecular mechanism of action is frequently far from obvious^5^. It is widely thought that these non-coding variants likely influence disease risk by functioning within *cis*-regulatory elements and altering expression of effector genes within their corresponding topologically associating domain (TAD). These effector genes are not necessarily the most proximal gene to the association signal, as *cis*-regulatory elements can influence gene expression up to megabases away. Therefore, functional characterization must be carried out to determine specifically which variants are causal and in turn which corresponding effector genes, near or far, confer susceptibility to disease. To date, most attention has been focused on only the very strongest GWAS loci, such as at *FTO*^26–28^, while many other significantly associated loci that rank lower in the signal list remain relatively understudied.

Our latest childhood obesity trans-ancestral GWAS meta-analysis on behalf of the Early Growth Genetics (EGG) consortium identified 19 loci that achieved genome-wide significance^22,23^. This included a locus on chr12q13 that was initially annotated based on its nearest gene, *FAIM2.* This signal has also been independently reported for obesity risk in children^29,30^ and adults^31–34^ across several ancestral populations. Crucially, this locus is more pronounced in children than in adults and ranks among more well-studied loci such as *FTO*, *MC4R*, *TMEM18*, and *BDNF* in the pediatric setting^23^; as such, it has been less studied to date given its less obvious role in adult obesity pathogenesis.

Our variant-to-gene mapping efforts have identified candidate *cis*-regulatory elements harboring childhood obesity risk variants in human embryonic stem cell (ESC)-derived hypothalamic neurons^12^ and other neural cell types^35^. In this study, we elected to characterize the relatively understudied chr12q13 locus and investigate the function of the lead candidate causal variant, rs7132908. We observed that the rs7132908 region contacts the promoters of *FAIM2* and several other genes within its TAD^12,35^ and therefore nominated these genes as candidate effector genes at this locus, with *FAIM2* having additional support via colocalization with expression quantitative trait loci (eQTL) data^36^. FAIM2 protects neurons from Fas-induced apoptosis^37,38^ and regulates neurite outgrowth^39^, neuroplasticity^40^, and synapse formation^41^ but has not been directly implicated in obesity pathogenesis to date. In this study, we initially used reporter assays in a relatively accessible astrocyte cell model to characterize the *cis*-regulatory activity of the region harboring rs7132908 and found that this variant regulated *FAIM2* expression with allele-specificity. Next, we used an established differentiation protocol^7^ to generate a relatively challenging model of hypothalamic neural progenitors and a heterogeneous population of hypothalamic neurons that were homozygous for either rs7132908 allele. We used bulk ATAC-seq pre-differentiation and single-nucleus ATAC-seq post-differentiation, when the cells were heterogenous, to assess chromatin accessibility both at rs7132908 and globally. We found that the rs7132908 region transitioned from closed to open chromatin during differentiation from ESCs to hypothalamic neurons. We also used bulk or single-nucleus RNA-seq to characterize changes in gene expression within the rs7132908 TAD and globally at three timepoints throughout differentiation, finding that rs7132908 genotype regulated the expression of *FAIM2* and other genes in multiple cell types. Finally, we report the striking observation that the rs7132908 obesity risk A allele decreased the proportion of neurons generated from our differentiation protocol from 61% to 11%. As such, we outline how our data strongly implicates rs7132908 as a causal variant at the chr12q13 obesity locus and nominates *FAIM2* as one candidate effector gene at this genomic location for further study.

## RESULTS

### rs7132908 is the putative causal variant at the chr12q13 childhood obesity locus

Each GWAS locus represents a genomic region harboring many single nucleotide polymorphisms (SNPs) in strong linkage disequilibrium, where any of these variants could be potentially causal and responsible for driving the significant association with the given trait. To implicate candidate causal non-coding variants at the chr12q13 obesity locus, our initial trans-ancestral fine mapping refined this specific signal to a 99% credible set consisting of six SNPs^23^. More recently, Bayesian fine mapping further refined this signal to 95% credible sets of 1-3 SNPs depending on which body weight trait definition was used^42^. These credible sets consistently implicate rs7132908 as the variant with the highest computed probability of being causal at this locus^23,42^. The childhood obesity risk A allele is common, with reported frequencies ranging from 10.25-60.53% across ethnicities^43^ and 28.86% globally^44^. In addition to childhood obesity, this locus is also associated with related traits such as increased body mass index (BMI) in adults, increased weight in adults, elevated type 2 diabetes susceptibility, and earlier age at menarche^45^.

After implicating a strong candidate causal variant computationally, we conducted a comprehensive characterization of the evidence supporting the *cis*-regulatory activity of the surrounding region in various cell types. First, we used our established variant-to-gene mapping approach which implicates potential *cis*-regulatory elements at GWAS loci using ATAC-seq to identify regions of accessible chromatin and then integrates high-resolution promoter-focused Capture-C or Hi-C to identify distal promoter interactions with those given open regions^12,35,46–49^. At the chr12q13 locus, we observed that rs7132908 resides within a putative *cis*-regulatory element in several human neural cell types, including primary astrocytes, induced pluripotent stem cell (iPSC)-derived cortical neural progenitors, ESC-derived hypothalamic neural progenitors, iPSC-derived cortical neurons, and ESC-derived hypothalamic neurons (**Fig. 1A, Supp. Table 1**)^12,35^. This is consistent with publicly available data from the Encyclopedia of DNA Elements (ENCODE) consortium’s ‘Registry of candidate *cis-*Regulatory Elements’ (version 3) which has annotated a cell type-agnostic candidate distal enhancer encompassing rs7132908 (candidate *cis-*regulatory element EH38E3015886)^50^. Second, we predicted the impact of the obesity risk A allele on transcription factor binding and identified 12 transcription factors that potentially regulate gene expression at the chr12q13 locus, namely HNF4A, HNF4G, PRD14, PRDM14, SRBP2, SREBF1, SREBF2, ZN143, ZN423, ZN554, ZN768, and ZNF416. Taken together, we concluded that rs7132908 is a strong candidate causal variant with predicted effects on gene expression through *cis*-regulatory mechanisms.

**Figure 1.**
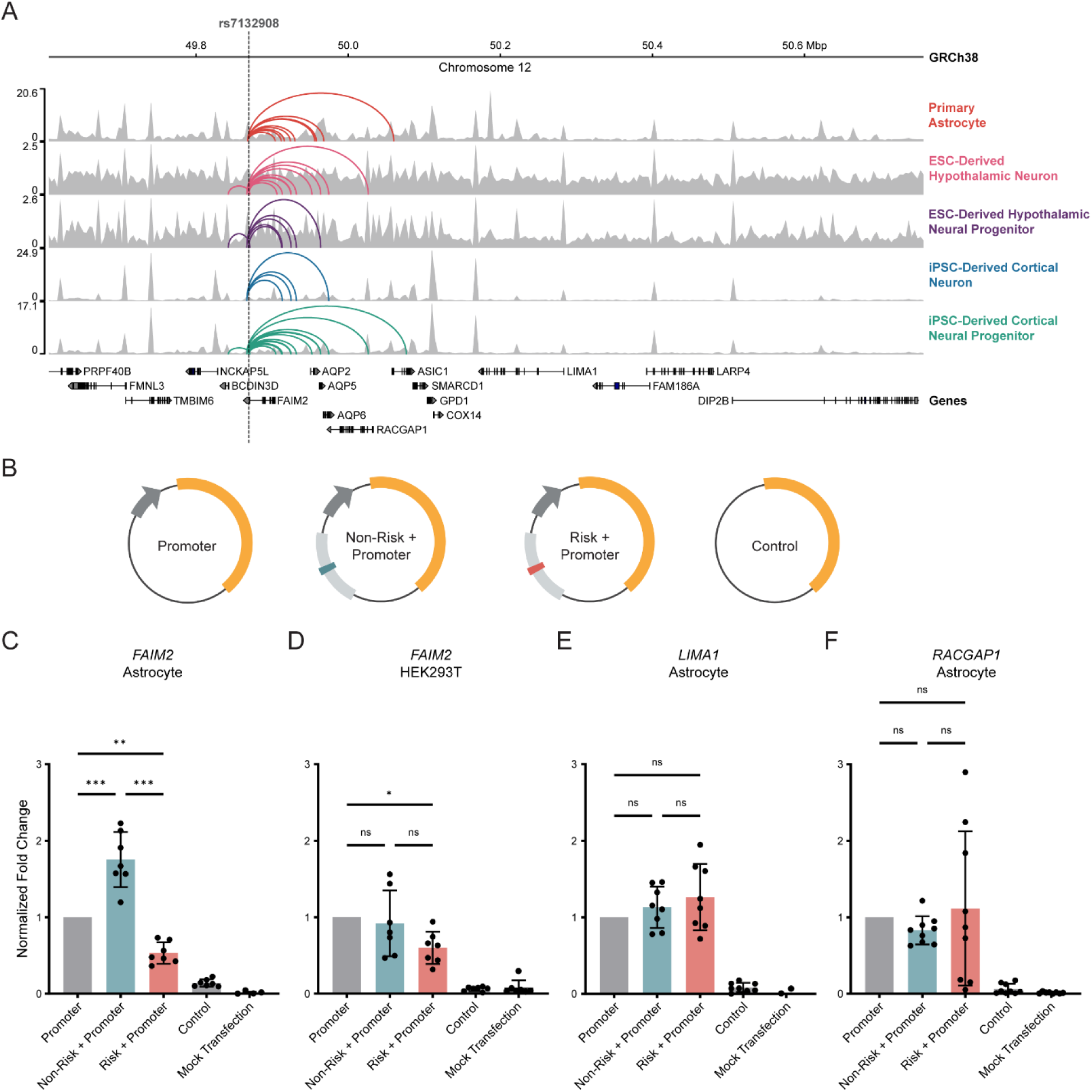
rs7132908 regulates FAIM2 expression with allele- and cell type-specificity. A) Chromatin accessibility represented by ATAC-seq tracks depicting normalized reads and chromatin loops at the TAD containing rs7132908 in neural cell types. Chromatin loops represent significant contacts between regions of open chromatin that harbored rs7132908 and a gene promoter. Grey dashed vertical line indicates rs7132908 position. B) Graphic representation of firefly luciferase reporter vectors used in luciferase reporter assays. C-F) Fold change of firefly luciferase fluorescence normalized to the promoter only control vector driven by the FAIM2 promoter in primary astrocytes (n = 7 biological replicates) (C), FAIM2 promoter in HEK293Ts (n = 7 biological replicates) (D), LIMA1 promoter in primary astrocytes (n = 8 biological replicates) (E), and RACGAP1 promoter in primary astrocytes (n = 9 biological replicates) (F). Data are represented as mean ± SD. *P-value < 0.05, **P-value < 0.01, ***P-value < 0.001 by one-way ANOVA with Tukey’s correction for multiple comparisons.

### *FAIM2* is the lead candidate effector gene at the chr12q13 childhood obesity locus

Chromosome conformation capture methods identify physical interactions between genomic regions and can nominate possible functional relationships, such as enhancer-promoter interactions. Analysis of our data indicated that the putative *cis*-regulatory element harboring rs7132908 interacted variably with the promoters of *AC025154.2*, *AQP2*, *AQP5*, *AQP6*, *ASIC1*, *BCDIN3D*, *FAIM2*, *LIMA1*, *LINC02395*, *LINC02396*, and *RACGAP1* across neural cell types (**Fig. 1A, Supp. Table 1**)^12,35^, suggesting potential temporal and cell type-specific control of multiple genes in the region, in a similar manner to the *FTO* locus^26,27^. We then performed colocalization analysis to intersect eQTL signals from the Genotype-Tissue Expression project (GTEx) with our variant-to-gene mapping results^36^. With the conservative overlap of the two approaches, we found only one gene at the chr12q13 locus, *FAIM2*, to be implicated by both our variant-to-gene mapping approach and eQTL analyses (**Supp. Table 1)**. We therefore implicated *FAIM2* as a primary candidate effector gene at this locus.

### Hypothalamic neurons and astrocytes are relevant *in vitro* models to study the regulatory effects of rs7132908 genotype

rs7132908 is located in the 3’ untranslated region (UTR) of *FAIM2* and 34,612 base pairs (bp) from the *FAIM2* transcription start site. The physical interaction between rs7132908 and the *FAIM2* promoter was observed in three neural cell types: primary astrocytes, iPSC-derived cortical neural progenitors, and ESC-derived hypothalamic neurons (**Fig. 1A, Supp. Table 1**)^12,35^. We measured gene expression in the neural cell types using bulk RNA-seq to aid in prioritizing *in vitro* models for our study. *FAIM2* expression was 2.26 transcripts per million (TPM) in iPSC-derived cortical neural progenitors, 42.85 TPM in primary astrocytes, and 136.75 TPM in ESC-derived hypothalamic neurons (**Supp. Table 1**)^12,35^. We have also previously identified that BMI-associated variants are significantly enriched in *cis*-regulatory elements annotated in an ESC-derived hypothalamic neuron model^12^. While this significant enrichment has not been detected in primary astrocytes, 7 out of 9 candidate effector genes nominated at the chr12q13 locus in ESC-derived hypothalamic neurons were also nominated in primary astrocytes (**Fig. 1A, Supp. Table 1**), suggesting similar genomic architecture in this region in these two cellular settings. Therefore, ESC-derived hypothalamic neurons and primary astrocytes were selected as *in vitro* models for studying the putative *cis*-regulatory relationship between rs7132908 and *FAIM2*, as well as other genes within their given TAD.

### rs7132908 regulates *FAIM2* expression with allele- and cell type-specificity

Many commonly used reporter assays to assess *cis*-regulatory function require an *in vitro* cell model that can be efficiently transfected. Neuron-like cells produced by stem cell differentiation are post-mitotic and transfection of these cells is very inefficient. For this reason, and given the comparable observations described above, we nominated primary astrocytes as a model to characterize the *cis*-regulatory function of the region harboring rs7132908 with luciferase reporter assays.

Given the ENCODE consortium’s ‘Registry of candidate *cis-*Regulatory Elements’ (version 1) annotated a cell type-agnostic regulatory element with a distal enhancer-like signature surrounding rs7132908 at chr12:49,868,837-49,869,798 (GRCh38)^50^, we elected to clone this putative enhancer sequence with an additional 50 bp flanking each side along with the *FAIM2* promoter sequence into a luciferase reporter vector. We then used site-directed mutagenesis to introduce the childhood obesity risk A allele at rs7132908. Each of these vectors, as well as normalization and negative control vectors (**Fig. 1B**), were transfected into astrocytes and luciferase fluorescence was quantified approximately 20 hours post-transfection.

We observed that the putative enhancer sequence with the non-risk allele significantly increased luciferase expression 1.75-fold when normalized to co-transfection and *FAIM2* promoter controls (adjusted *P*-value < 0.001) (**Fig. 1C**). In contrast, the same vector with a single base change to the obesity risk A allele significantly decreased luciferase expression 0.53-fold after normalization (adjusted *P*-value = 0.003) (**Fig. 1C**). We then sought to carry out this experiment using HEK293T cells to determine if this *cis-*regulatory activity could also be observed in a non-neural cell type. In HEK293Ts, we found that the putative enhancer sequence harboring the non-risk G allele did not significantly increase luciferase expression, while the obesity risk allele decreased luciferase expression by 0.60-fold after normalization (adjusted *P*-value = 0.037) (**Fig. 1D**). From this, we conclude that the rs7132908 obesity risk allele negatively regulates expression from the *FAIM2* promoter in astrocytes but displays weaker effects in non-neuronal HEK293Ts.

In addition to *FAIM2,* our 3D genomic variant-to-gene mapping efforts in primary astrocytes also nominated *LIMA1* and *RACGAP1* as possible effector genes at the chr12q13 locus (**Fig. 1A, Supp. Table 1**). This was determined using the criteria that the promoters of these genes interacted with rs7132908, the promoters of these genes and rs7132908 were both in open chromatin, and that these genes were expressed (TPM > 1) in this cell type. However, when we assessed the *cis-*regulatory activity of this region with the *LIMA1* and *RACGAP1* promoter sequences in astrocytes, we observed no significant changes in luciferase expression with either rs7132908 allele after normalization, although we note the results for the risk A allele with the *RACGAP1* promoter were highly variable (**Fig. 1E-F**).

### The putative *cis*-regulatory region harboring rs7132908 is inactive in ESCs

After characterizing the *cis*-regulatory activity of the region harboring rs7132908 in astrocytes, we were motivated to characterize the effect of the rs7132908 childhood obesity risk allele in cells at multiple timepoints throughout differentiation to hypothalamic neurons. We used the H9 ESC line, which is homozygous for the rs7132908 non-risk G allele, and leveraged CRISPR-Cas9 homology-directed repair to generate three isogenic, clonal lines that were homozygous for the rs7132908 risk A allele.

To characterize chromatin accessibility in homogenous ESCs, we performed bulk ATAC-seq using three replicates of the parent ESC line and the three clonal ESC lines generated with CRISPR. We detected 240,760 peaks but then performed filtering to limit our analysis to peaks present in at least two samples and removed peaks with low read support (one count per million or less), which reduced our peak list to 94,543. We observed that the first principal component, explaining 27.8% of the variation between samples, was due to genotype at rs7132908 (**Fig. 2A**). 286 peaks were differentially accessible using an adjusted *P*-value < 0.05 and |log2 fold change| > 1 (**Fig. 2B, Supp. Table 2**). 145 peaks were significantly more accessible in the ESCs with the non-risk allele and 141 peaks were more accessible with the risk allele. However, rs7132908 itself was not found in a peak of accessible chromatin in these undifferentiated ESCs (**Fig. 2C**).

**Figure 2.**
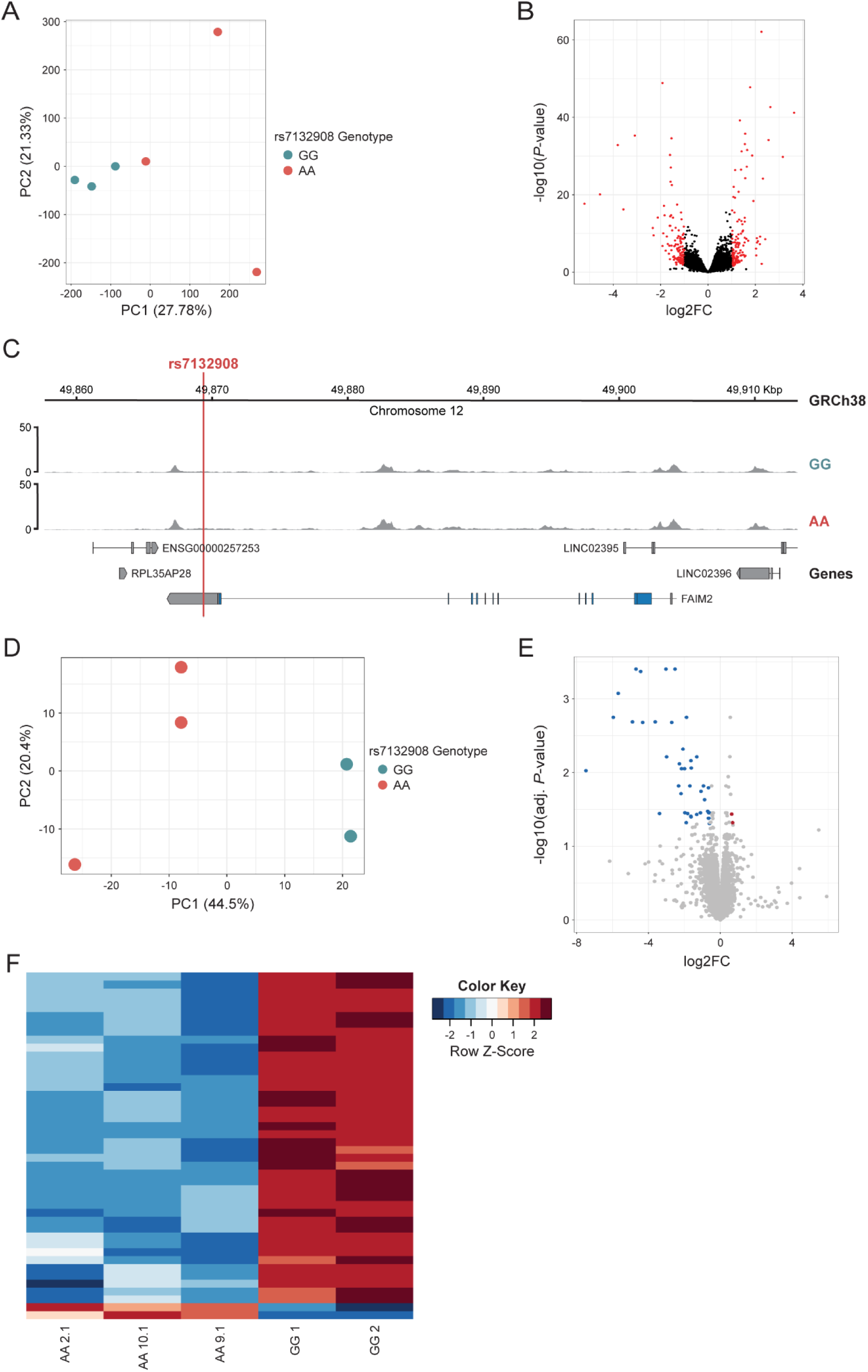
The putative cis-regulatory region harboring rs7132908 is inactive in ESCs. A) PCA plot of ESC ATAC-seq libraries (GG n = 3, AA n = 3 lines). B) Volcano plot of adjusted P-values (-log10) and fold change (log2) of a total of 94,543 ATAC-seq peaks tested for differential accessibility due to the rs7132908 obesity risk allele in ESCs. Red dots indicate significantly differentially accessible peaks (adjusted P-value < 0.05 and |log2 fold change| > 1) in ESCs homozygous for the obesity risk allele and black dots indicate peaks with no significant differences in accessibility. C) Chromatin accessibility represented by ATAC-seq tracks depicting normalized reads across FAIM2 in ESCs homozygous for either rs7132908 allele. Red vertical line indicates rs7132908 position. D) PCA plot of ESC RNA-seq libraries (GG n = 2, AA n = 3 lines). E) Volcano plot of adjusted P-values (-log10) and fold change (log2) of a total of 14,581 genes tested for differential expression due to the rs7132908 obesity risk allele in ESCs. Blue dots indicate significantly down-regulated genes (adjusted P-value < 0.05 and log2 fold change < -0.58) and red dots indicate significantly up-regulated genes (adjusted P-value < 0.05 and log2 fold change > 0.58) in ESCs homozygous for the obesity risk allele. Grey dots indicate genes with no significant differences in expression. F) Heatmap depicting significantly differentially expressed genes (adjusted P-value < 0.05 and |log2 fold change| > 0.58) due to the rs7132908 obesity risk allele in ESCs.

To identify any transcriptional differences due to rs7132908 genotype in homogenous ESCs, we performed bulk RNA-seq using two replicates of the parent ESC line and the three clonal ESC lines generated with CRISPR. We achieved reads from 35,595 genes and then performed filtering to detect those expressed at greater than one count per million in at least two samples, which reduced our gene list to 14,581. We observed that the first principal component, explaining 44.5% of the variation between samples, was due to genotype at rs7132908 (**Fig. 2D**). 44 genes were differentially expressed using an adjusted *P*-value < 0.05 and |log2 fold change| > 0.58 (**Fig. 2E-F, Supp. Table 3**). 42 genes were significantly down-regulated in the rs7132908 risk A allele homozygote ESCs, while just two genes were up-regulated. As most enhancer-promoter interactions are known to occur within the same TAD, we were motivated to determine if rs7132908 genotype affected the expression of genes within its TAD. However, none of the genes in the TAD harboring rs7132908^51^ were differentially expressed in this undifferentiated ESC setting. Taken together, we observed relatively small changes in expression and accessibility due to the introduction of the obesity risk allele in undifferentiated ESCs, consistent with the notion that rs7132908 primarily functions in neural cells.

### rs7132908 genotype influences gene expression in ESC-derived hypothalamic neural progenitors

To generate hypothalamic neural progenitors and characterize the effects of rs7132908 at this stage, we differentiated two replicates of the parent ESC line which were homozygous for the rs7132908 non-risk G allele and the two clonal ESC lines generated with CRISPR which were homozygous for the rs7132908 risk A allele for 14 days using an established protocol (**Fig. 3A**)^7^. Day 14 was selected given it was the last day after direction towards ventral diencephalon hypothalamic identity and cell cycle exit but before neuron maturation^7^. We compared the global transcriptomic profile of the hypothalamic neural progenitors homozygous for the rs7132908 non-risk allele to profiles of primary human tissues in the GTEx RNA-seq database^36^ (donor ages 20-71 years old, with 68.1% 50 years or older) as well as primary human pediatric hypothalamus tissue from three donors homozygous for the rs7132908 non-risk allele (donor ages 4-14 years old, average age = 8.67). The non-risk hypothalamic neural progenitors most highly correlated with the primary human pediatric hypothalamus tissue (correlation coefficient = 0.80, *P*-value < 0.001) (**Fig. S1A**).

**Figure 3.**
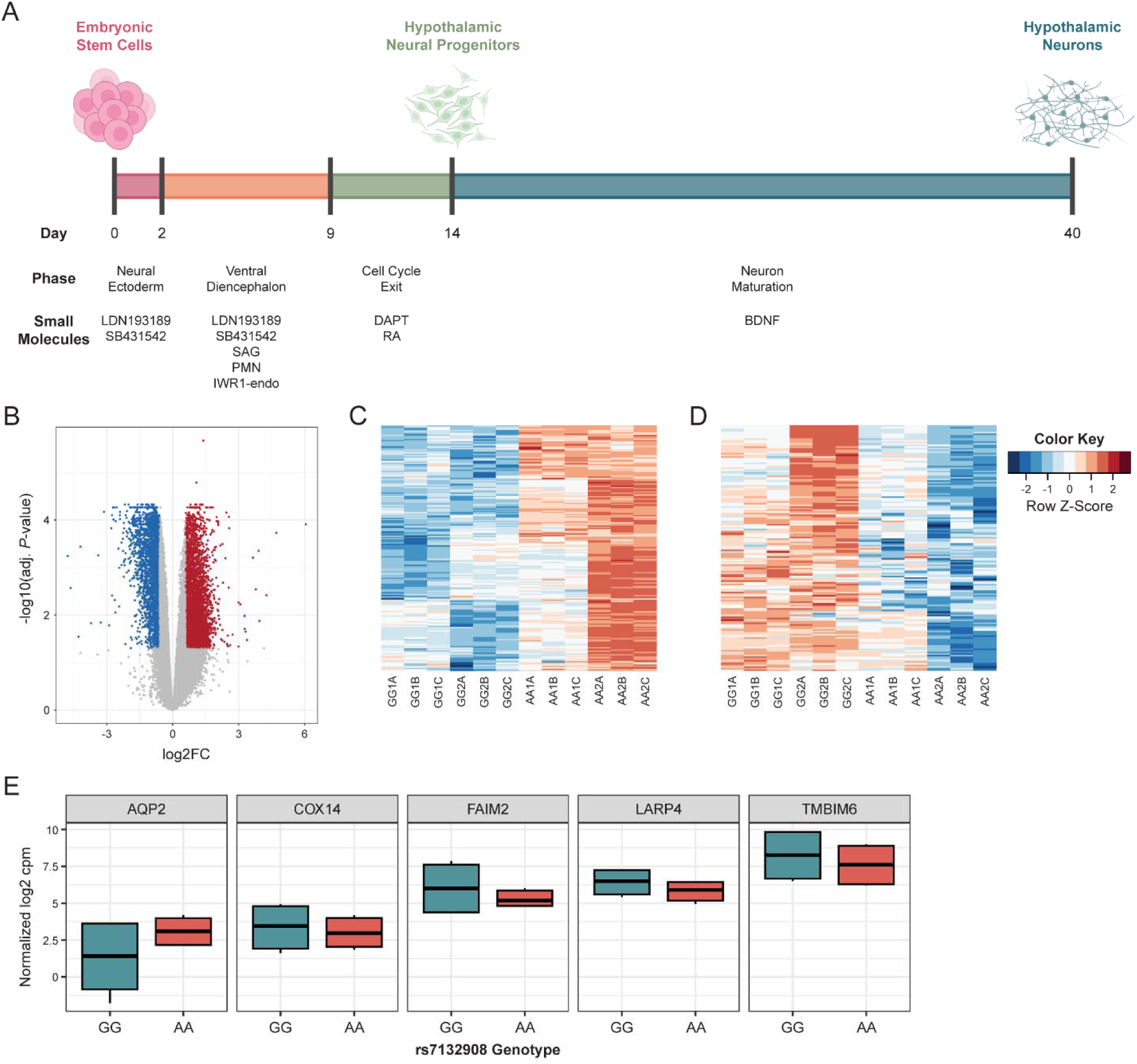
rs7132908 genotype influences gene expression in ESC-derived hypothalamic neural progenitors. A) Schematic of differentiation of ESCs to hypothalamic neurons, including duration, phases, and key small molecules to direct cell fates. B) Volcano plot of adjusted P-values (-log10) and fold change (log2) of a total of 29,302 genes tested for differential expression due to the rs7132908 obesity risk allele in hypothalamic neural progenitors. Blue dots indicate significantly down-regulated genes (adjusted P-value < 0.05 and log2 fold change < -0.58) and red dots indicate significantly up-regulated genes (adjusted P-value < 0.05 and log2 fold change > 0.58) in hypothalamic neural progenitors homozygous for the obesity risk allele. Grey dots indicate genes with no significant differences in expression. C) Heatmap depicting module 4 genes significantly up-regulated (adjusted P-value < 0.05 and log2 fold change > 0.58) due to the rs7132908 obesity risk allele in hypothalamic neural progenitors. D) Heatmap depicting module 5 genes significantly down-regulated (adjusted P-value < 0.05 and log2 fold change < -0.58) due to the rs7132908 obesity risk allele in hypothalamic neural progenitors. E) Box plots of gene expression (normalized log2 cpm) for genes in the rs7132908 TAD that were significantly differentially expressed (adjusted P-value < 0.05, |log2 fold change| > 0.58). See also Figure S1.

**Supplemental Figure 1.**
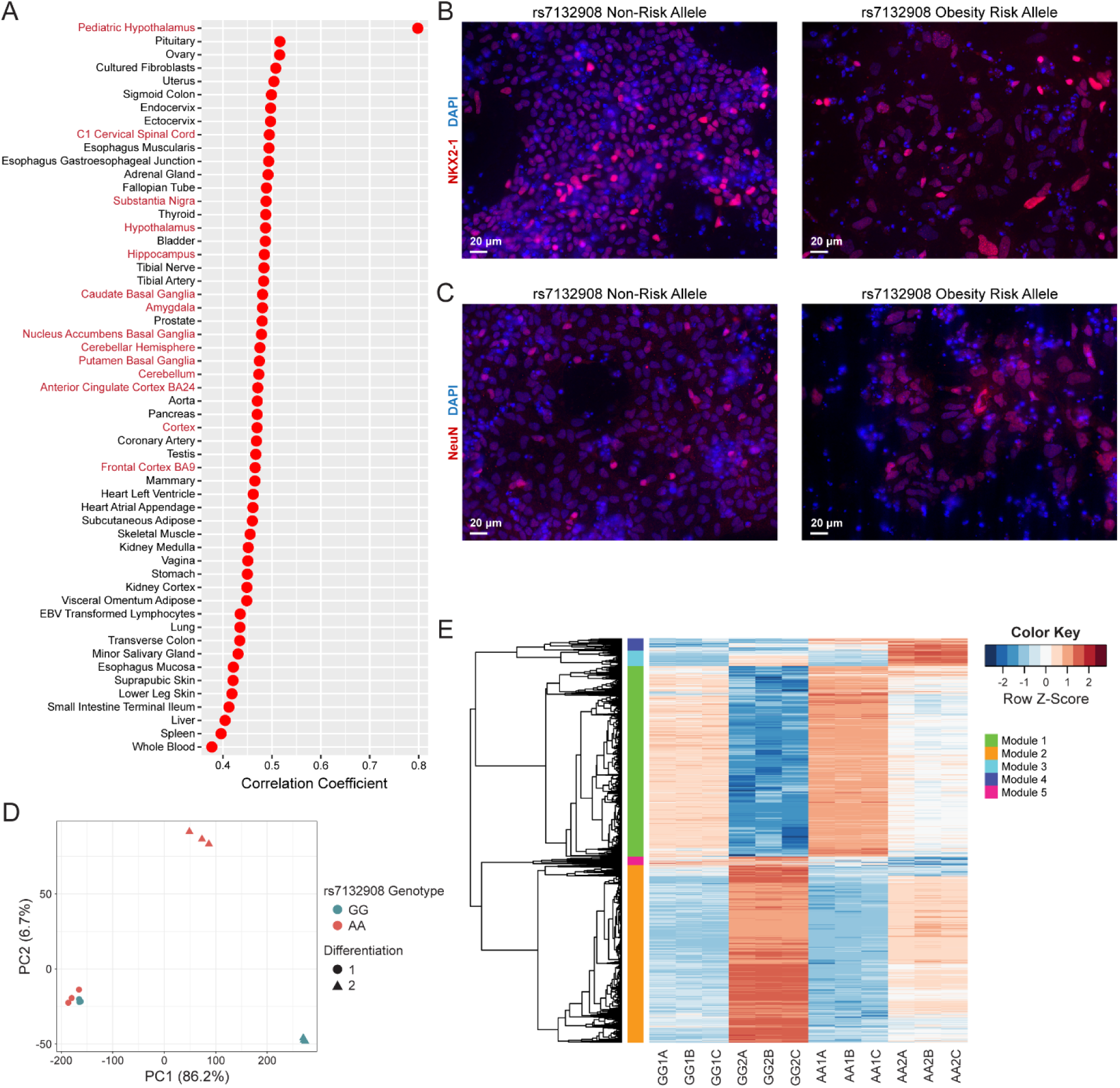
Hypothalamic neural progenitors, related to Figure 3. A) Dot plot of Spearman’s rank correlation coefficients resulting from comparing TPMs of 16,159 genes expressed in rs7132908 non-risk allele homozygous hypothalamic neural progenitors, rs7132908 non-risk allele homozygous human pediatric hypothalamus tissue, and human tissues or cells in the GTEx RNA-seq database. Red dots indicate significant correlations (P-value < 0.05). Tissue names in red indicate brain tissues. B-C) Representative images of hypothalamic neural progenitors on day 14, with immunostaining for a marker of the developing hypothalamus, NKX2-1 (red) (B) and a marker of post-mitotic neurons, NeuN (red) (C) (scale bar = 20 µm). Nuclei were stained with DAPI (blue). Cells were homozygous for either the rs7132908 non-risk allele (left) or obesity risk allele (right). D) PCA plot of hypothalamic neural progenitor RNA-seq libraries (GG n = 2 biological replicates with 3 technical replicates each, AA n = 2 biological replicates with 3 technical replicates each). E) Heatmap depicting significantly differentially expressed genes (adjusted P-value < 0.05, |log2 fold change| > 0.58) due to the rs7132908 obesity risk allele in hypothalamic neural progenitors. Genes were clustered into 5 modules using hierarchical clustering (green, orange, light blue, dark blue, pink).

To identify transcriptional differences due to rs7132908 genotype in homogeneous hypothalamic neural progenitors, we performed bulk RNA-seq. We detected reads mapping to 35,595 genes and then performed filtering to detect those expressed at greater than one count per million in at least two samples which reduced our gene list to 29,302. We observed that the first principal component, explaining 86.2% of the variation between samples, was due to batch as we differentiated pairs of non-risk and risk allele cells at two separate times (**Fig. S1D**). Therefore, we incorporated batch information as a covariate in our linear model to adjust for this effect for our differential expression analysis. As a result, 6,494 genes were differentially expressed using an adjusted *P*-value < 0.05 and |log2 fold change| > 0.58 (**Fig. 3B, Supp. Table 4**). Of these, 3,232 genes were significantly down-regulated in the neural progenitors homozygous for the rs7132908 risk allele, while 3,262 genes were up-regulated. Five genes in the TAD harboring rs7132908^51^ were differentially expressed. *FAIM2* and three other genes (*TMBIM6*, *LARP4*, and *COX14)* were down-regulated in these neural progenitors homozygous for the rs7132908 risk A allele and *AQP2* was up-regulated (**Fig. 3E**).

To explore global changes in gene expression, we clustered the differentially expressed genes into five modules with hierarchical clustering (**Fig. S1E**) and selected two modules (modules 4 and 5) representing the genes most strongly differentially expressed due to genotype at rs7132908 for downstream analysis. Module 4 consisted of 216 genes consistently up-regulated in neural progenitors homozygous for the rs7132908 risk A allele (**Fig. 3C**). Functional enrichment analysis of the module 4 up-regulated genes identified significantly enriched Gene Ontology terms^52,53^, including biological processes such as programmed cell death, apoptotic process, and intrinsic apoptotic signaling pathway in response to endoplasmic reticulum stress (**Supp. Table 5**). Module 5 consisted of 152 genes consistently down-regulated in neural progenitors homozygous for the rs7132908 risk allele (**Fig. 3D**). The module 5 down-regulated genes were also used to determine any enriched Gene Ontology terms^52,53^, however, no significantly enriched biological processes were identified (**Supp. Table 5**).

### ESC-derived hypothalamic neurons molecularly resemble the human hypothalamus

Next, we generated hypothalamic-like neurons using two replicates of the parent ESC line which were homozygous for the rs7132908 non-risk G allele and the two clonal ESC lines generated with CRISPR which were homozygous for the rs7132908 risk A allele for 40 days in four independent differentiations using an established protocol^7^ and then collected nuclei (**Fig. 3A**). Day 40 was selected given a previous characterization of this protocol found that this duration was sufficient to produce heterogenous populations of functional neurons that closely resemble those found in the human hypothalamus^7^. These nuclei from several cell types were used to simultaneously profile gene expression and open chromatin in each cell using a multi-omic single-nucleus RNA-seq and ATAC-seq approach.

After quality control, a previously published human hypothalamus single-cell RNA-seq reference dataset^54^ was used to identify cell types in our dataset (**Fig. S2D**). To ensure that the cell type identifications were likely to be accurate, we prioritized cells with high-confidence annotations using a classification score threshold (≥ 0.8) that was previously demonstrated to increase accuracy^55^. This method identified cells annotated as neurons, oligodendrocyte precursors (OPCs), or fibroblasts based on their transcriptional profile with classification scores above our threshold (**Fig. 4A**). These annotations are further supported by expression patterns of known marker genes for each cell type, including *MAP2* and *TUBB3* for neurons and *COL1A1, COL1A2,* and *COL6A2* for fibroblasts (**Fig. 4B**). We note that the OPC population did not highly or uniformly express conventional marker genes, such as *PDGFRA, CSPG4, OLIG1, OLIG2,* and *SOX10* (**Fig. S2E**), although this population did express cell cycle genes, such as *CENPF* and *TOP2A*, which have been observed in OPCs^56^ and neural intermediate progenitors^57^ (**Fig. 4B**).

**Figure 4.**
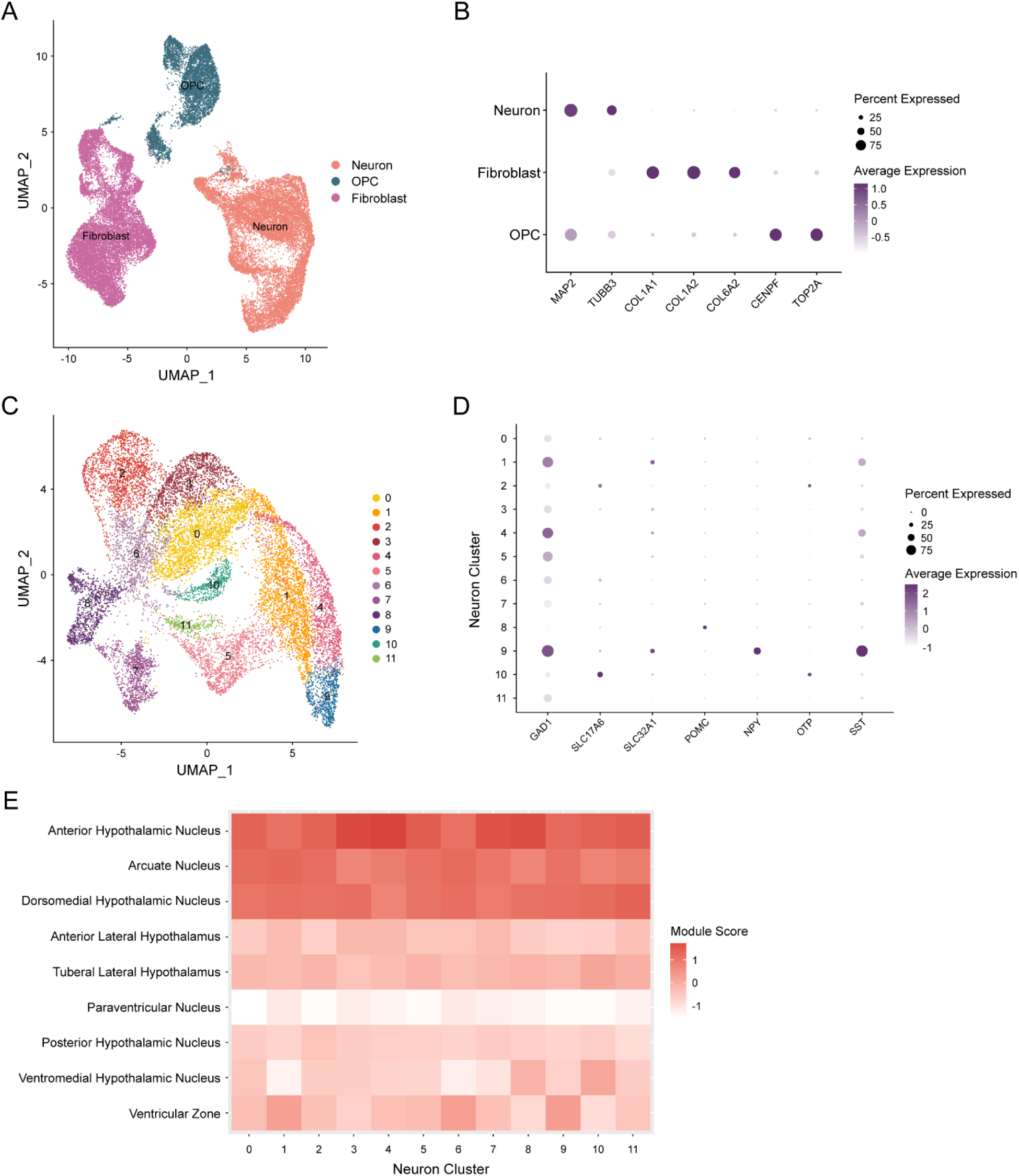
ESC-derived hypothalamic neurons molecularly resemble the human hypothalamus. A) UMAP depicting all cells clustered by single-nucleus RNA-seq profile and annotated by cell type. B) Dot plot depicting average expression (scaled and log2 normalized counts) and percent of cells that expressed neuron (MAP2 and TUBB3), fibroblast (COL1A1, COL1A2, and COL6A2), and OPC (CENPF and TOP2A) marker genes, split by cell type. C) UMAP depicting all neurons clustered by single-nucleus RNA-seq profile and annotated by cluster identity. D) Dot plot depicting average expression (scaled and log2 normalized counts) and percent of cells that expressed inhibitory (GAD1), excitatory (SLC17A6), GABAergic (SLC32A1), and hypothalamic (POMC, NPY, OTP, and SST) neuron marker genes, split by cluster identity. E) Heatmap showing average module scores across all neuron clusters for each human prenatal hypothalamic nucleus gene set published in the Allen Brain Atlas database, plotted as the column Z-score per neuron cluster. See also Figure S2.

**Supplemental Figure 2.**
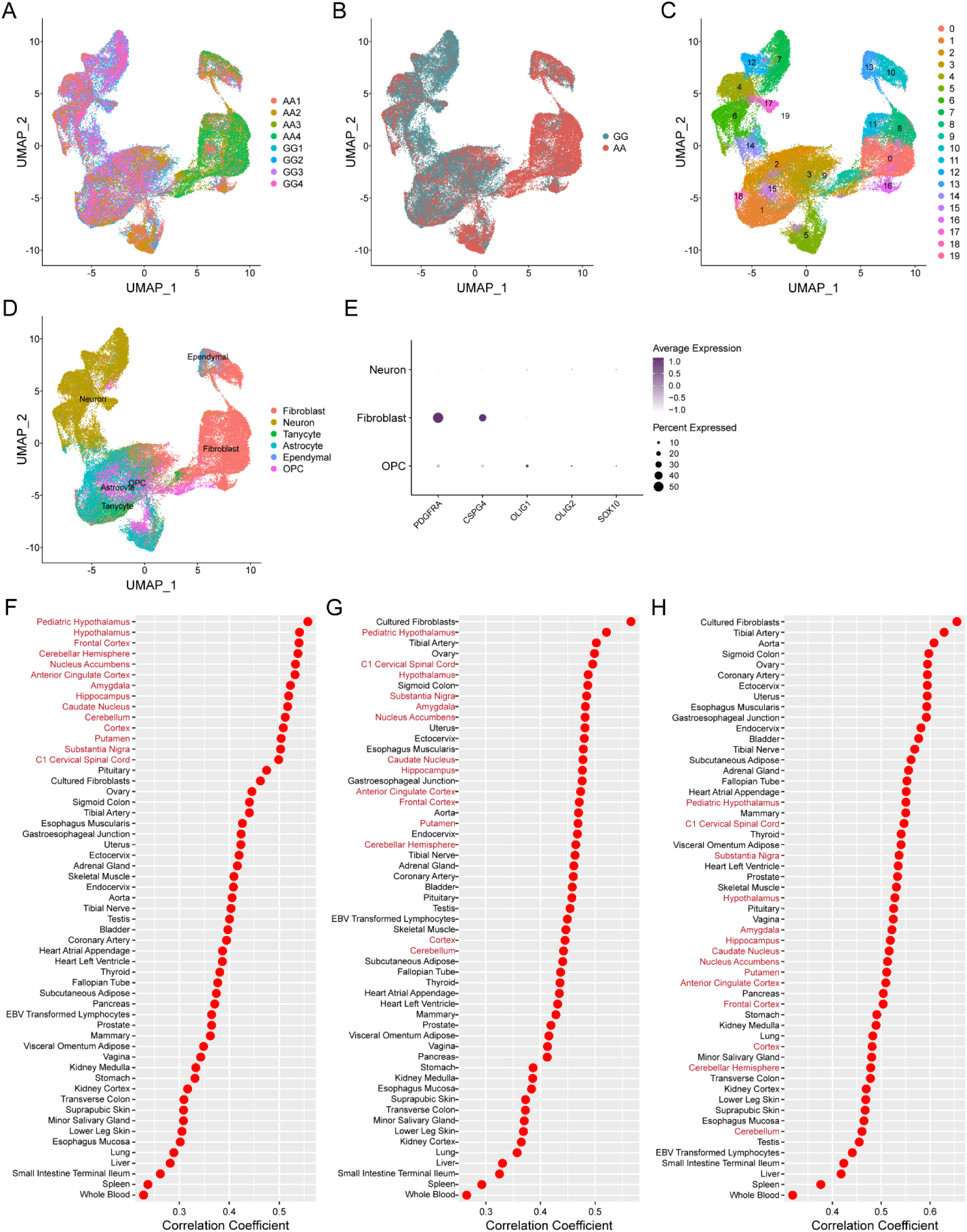
Hypothalamic single-nucleus RNA-seq analysis, related to Figure 4. **A-D)** UMAP depicting all cells clustered by single-nucleus RNA-seq profile and annotated by replicate sample (A), rs7132908 genotype (B), cluster identity (C), and predicted cell type annotation before cells with classification scores below the 0.8 threshold were removed (D). E) Dot plot depicting average expression (scaled and log2 normalized counts) and percent of cells that expressed canonical OPC marker genes (PDGFRA, CSPG4, OLIG1, OLIG2, and SOX10), split by cell type. F-H) Dot plots of Spearman’s rank correlation coefficients resulting from comparing TPMs of genes expressed in rs7132908 non-risk allele homozygous hypothalamic neurons (F), OPCs (G), and fibroblasts (H) to human pediatric hypothalamus tissue from donors homozygous for the rs7132908 non-risk allele and human tissues or cells in the GTEx RNA-seq database. Red dots indicate significant correlations (P-value < 0.05). Tissue names in red indicate brain tissues.

Additionally, we compared the transcriptomic signatures of each cell type to expression data in the GTEx RNA-seq database^36^ (donor ages 20-71 years old, with 68.1% 50 years or older) as well as primary human pediatric hypothalamus tissue from three donors homozygous for the rs7132908 non-risk allele (donor ages 4-14 years old, average age = 8.67). We found that the neurons were most strongly correlated with pediatric hypothalamus and adult hypothalamus (correlation coefficients = 0.56 and 0.54, respectively, *P*-values < 0.001), the OPCs correlated most strongly with fibroblasts and pediatric hypothalamus (correlation coefficients = 0.57 and 0.52, respectively, *P*-values < 0.001), and the fibroblasts most strongly correlated with fibroblasts and tibial artery (correlation coefficients = 0.66 and 0.63, respectively, *P*-values < 0.001) (**Fig. S2F-H**).

Within the neuron population (**Fig. 4C**), there were distinct expression patterns of markers for several neuron types, including inhibitory (*GAD1*), excitatory (*SLC17A6*), and GABAergic (*SLC32A1*) neurons (**Fig. 4D**). We also identified neuronal clusters expressing known hypothalamus genes, such as *POMC*, *NPY*, *OTP,* and *SST* (**Fig. 4D**). Next, we were motivated to compare the transcriptomic signatures of each neuronal cluster (**Fig. 4C**) to human prenatal hypothalamic subregion gene sets published in the Allen Brain Atlas database^58–61^, given that the neuron population displayed expression patterns most similar to pediatric hypothalamus tissue. We found that each cluster closely resembled the hypothalamic arcuate nucleus which regulates feeding behavior and energy expenditure^62^, the dorsomedial hypothalamic nucleus which regulates food intake and body weight^63^, and the anterior hypothalamic nucleus which regulates defensive behaviors^64^ (**Fig. 4E**).

### The putative *cis*-regulatory region harboring rs7132908 is active in ESC-derived hypothalamic cell types

We used the single-nucleus ATAC-seq data to characterize chromatin accessibility in the heterogenous ESC-derived hypothalamic cells. Unlike in the ESCs, the *cis*-regulatory element containing rs7132908 was open in all derived cell types (**Fig. 5A**). When comparing chromatin accessibility globally between rs7132908 genotypes across all annotated cells, 12,586 ATAC-seq peaks were differentially accessible using an adjusted *P*-value < 0.05 and |log2 fold change| > 1, with 5,604 peaks displaying decreased accessibility with the risk A allele and 6,982 peaks displaying increased accessibility (**Fig. 5B, Supp. Table 6**). We also detected transcription factor motifs enriched in differentially accessible peaks with an adjusted *P*-value < 0.005 and |log2 fold change| ≥ 1. We found that 565 transcription factor motifs were significantly enriched (adjusted *P*-value < 0.05) in peaks more accessible with the rs7132908 non-risk G allele and 446 were enriched in peaks more accessible with the risk A allele (**Supp. Table 7**). The peak harboring rs7132908 at chr12:49,868,731-49,869,775 (GRCh38) displayed decreased accessibility with the risk A allele by 27.62% (adjusted *P*-value = 1.08×10^-88^) when considering all annotated cells. We also repeated these analyses in each annotated cell type and detected 3,406, 12,386, and 7,543 significantly differentially accessible regions in neurons, OPCs, and fibroblasts, respectively (**Fig. 5C-E, Supp. Table 6**). The peak surrounding rs7132908 was less accessible with the risk A allele by 40.74% in fibroblasts (adjusted *P*-value = 1.35×10^-14^), but more accessible in neurons with the risk A allele by 78.92% (adjusted *P*-value = 2.31×10^-21^) and not significantly different in OPCs. We then identified significantly differentially accessible regions that were consistent between analyses when considering each individual cell type and all annotated cells combined (**Fig. 5F**) and their top enriched transcription factor motifs (**Fig. 5G**). We conclude that rs7132908 is in an active chromatin region post-differentiation and that the rs7132908 risk A allele influences accessibility both locally and globally.

**Figure 5.**
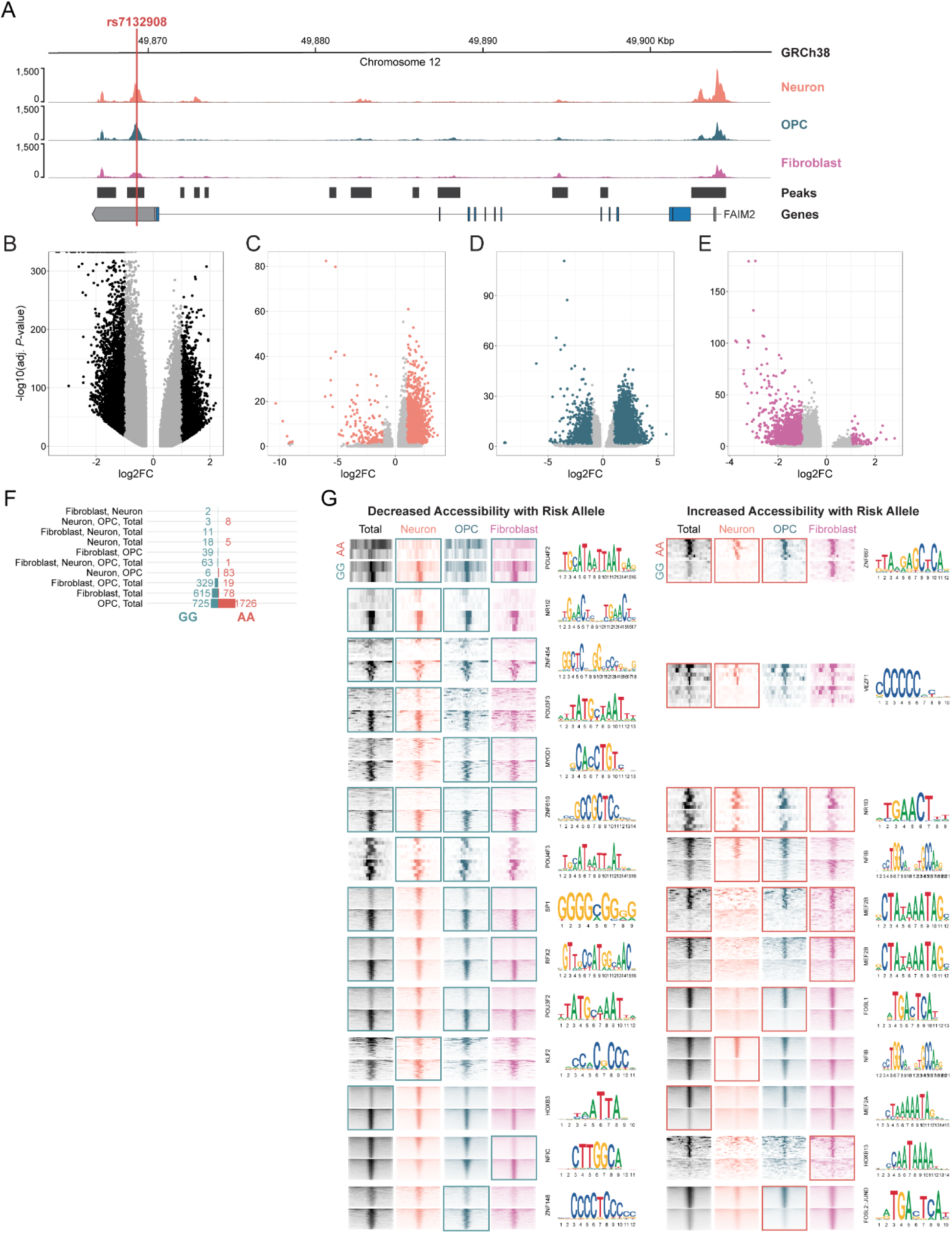
The putative cis-regulatory region harboring rs7132908 is active in ESC-derived hypothalamic cell types. A) Chromatin accessibility represented by ATAC-seq tracks depicting normalized reads across FAIM2 in ESC-derived neurons, OPCs, and fibroblasts. Red vertical line indicates rs7132908 position. B-E) Volcano plots of adjusted P-values (-log10) and fold change (log2) of ATAC-seq peaks tested for differential accessibility due to the rs7132908 obesity risk allele in total cells (B), neurons (C), OPCs (D), and fibroblasts (E). Black or colored dots indicate significantly differentially accessible peaks (adjusted P-value < 0.05 and |log2 fold change| > 1) and grey dots indicate peaks with no significant differences in accessibility. F) Bar plot of numbers of differentially accessible regions from B-E that overlapped between analyses. G) ATAC-seq read enrichment heatmaps for groups of regions categorized in F and their corresponding top-most enriched transcription factor binding motifs. Windows indicate which cell type(s) yielded such groups of differentially accessible regions.

### The rs7132908 obesity risk allele dramatically decreases the proportion of neurons produced by hypothalamic neuron differentiation

As expected, during each hypothalamic neuron differentiation, we began to observe neuron morphology with brightfield microscopy once the cells were exposed to BDNF in the neuron maturation phase (days 14-40) (**Fig. 3A**). Strikingly, there were fewer cells exhibiting neuron morphology for those homozygous for the rs7132908 risk A allele (**Fig. 6A**). To confirm this observation, we stained day 40 cells from each genotype to detect MAP2, which is a marker of mature neuron dendrites. Indeed, although each well was seeded at the same density and cultured simultaneously, fewer MAP2+ cells were observed in the risk A allele condition (**Fig. 6B**).

**Figure 6.**
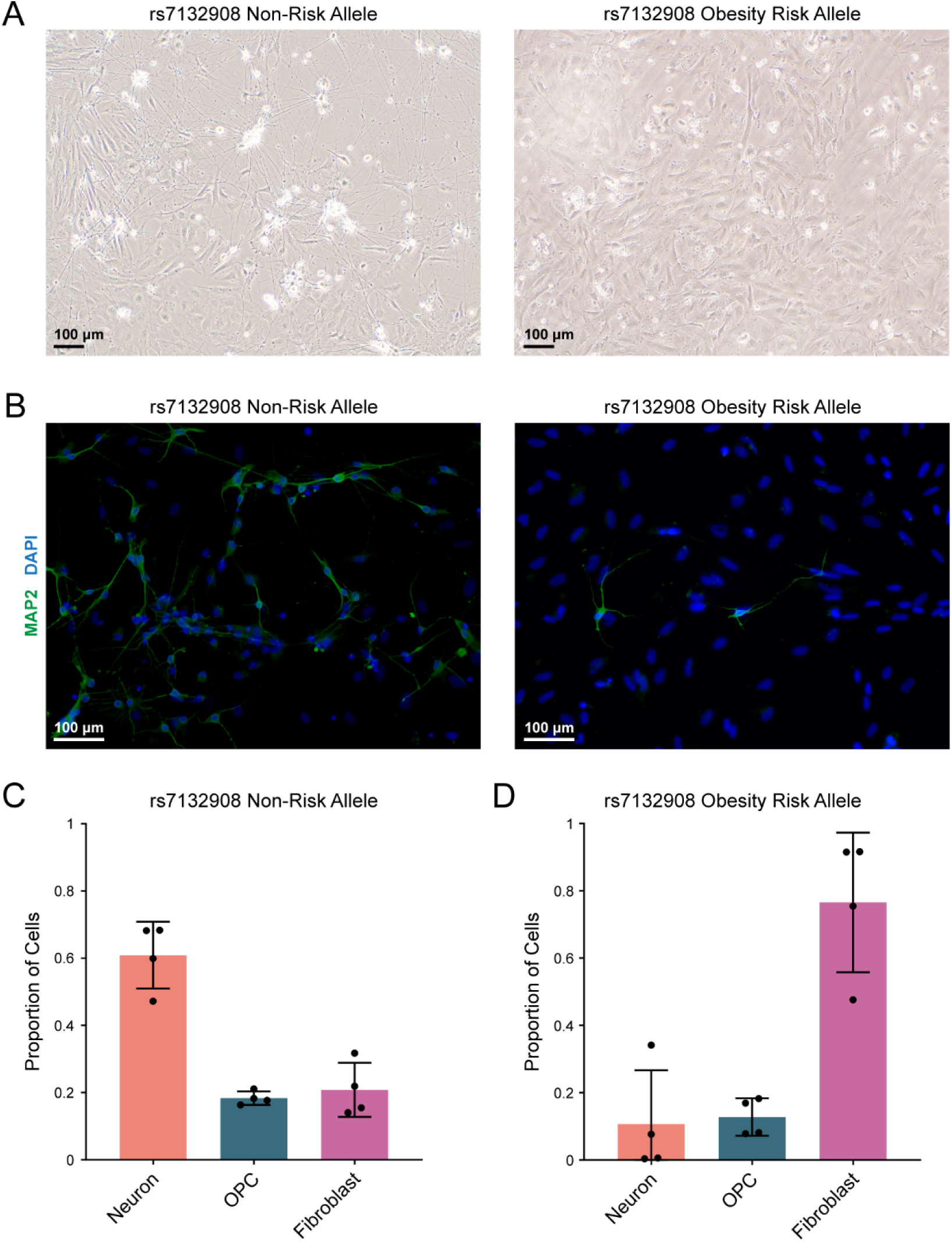
The rs7132908 obesity risk allele dramatically decreases the proportion of neurons produced by hypothalamic neuron differentiation. A) Representative brightfield images of hypothalamic neurons mid-differentiation on day 29 (scale bar = 100 µm). Cells were homozygous for either the rs7132908 non-risk allele (left) or obesity risk allele (right). B) Representative images of hypothalamic neurons post-differentiation on day 40, with immunostaining for a mature neuron marker, MAP2 (green) (scale bar = 100 µm). Nuclei were stained with DAPI (blue). Cells were homozygous for either the rs7132908 non-risk allele (left) or obesity risk allele (right). C-D) Proportion of total cells homozygous for the rs7132908 non-risk allele annotated as each cell type (n = 4 differentiation replicates) (C) and homozygous for the rs7132908 obesity risk allele annotated as each cell type (n = 4 differentiation replicates) (D). Data are represented as mean ± SD.

Next, we set out to further confirm this result using our annotated single-nucleus RNA-seq dataset. We partitioned the 38,044 annotated cells by genotype at rs7132908 and differentiation replicate sample, then quantified the proportions of cells identified as neurons, OPCs, or fibroblasts in each condition. On average, the cells homozygous for the rs7132908 non-risk G allele were comprised of 60.90% neurons, 18.33% OPCs, and 20.77% fibroblasts (**Fig. 6C**). In contrast, the cells homozygous for the rs7132908 risk A allele comprised of 10.69% neurons, 12.78% OPCs, and 76.53% fibroblasts (**Fig. 6D**). Taken together, we observed that a single base change from the rs7132908 non-risk G allele to the obesity risk A allele in the same genetic background is sufficient to substantially decrease the proportion of neurons produced by hypothalamic neuron differentiation.

### rs7132908 genotype influences gene expression in ESC-derived hypothalamic cell types

In addition to identifying differences in cell type proportions, we were also motivated to identify changes in gene expression due to genotype at rs7132908 in the ESC-derived hypothalamic cells. First, we included all cells that passed our quality control, detected 36,601 genes, and performed principal component analysis to determine that 85% of the variation between replicate samples was explained by rs7132908 genotype (**Fig. S3A**). We then identified that 6,409 genes were differentially expressed using an adjusted *P*-value < 0.05 and |log2 fold change| > 0.58 (**Fig. 7A**, **Fig. 7E, Supp. Table 8**). 3,212 genes were significantly down-regulated in the cells homozygous for the rs7132908 risk allele, while 3,197genes were up-regulated. Four genes in the TAD harboring rs7132908^51^, including *FAIM2*, were differentially expressed; two were down-regulated in cells homozygous for the rs7132908 risk A allele (*FAIM2,* and *ASIC1*) and two were up-regulated (*FMNL3* and *LIMA1*) (**Fig. 7I**).

**Figure 7.**
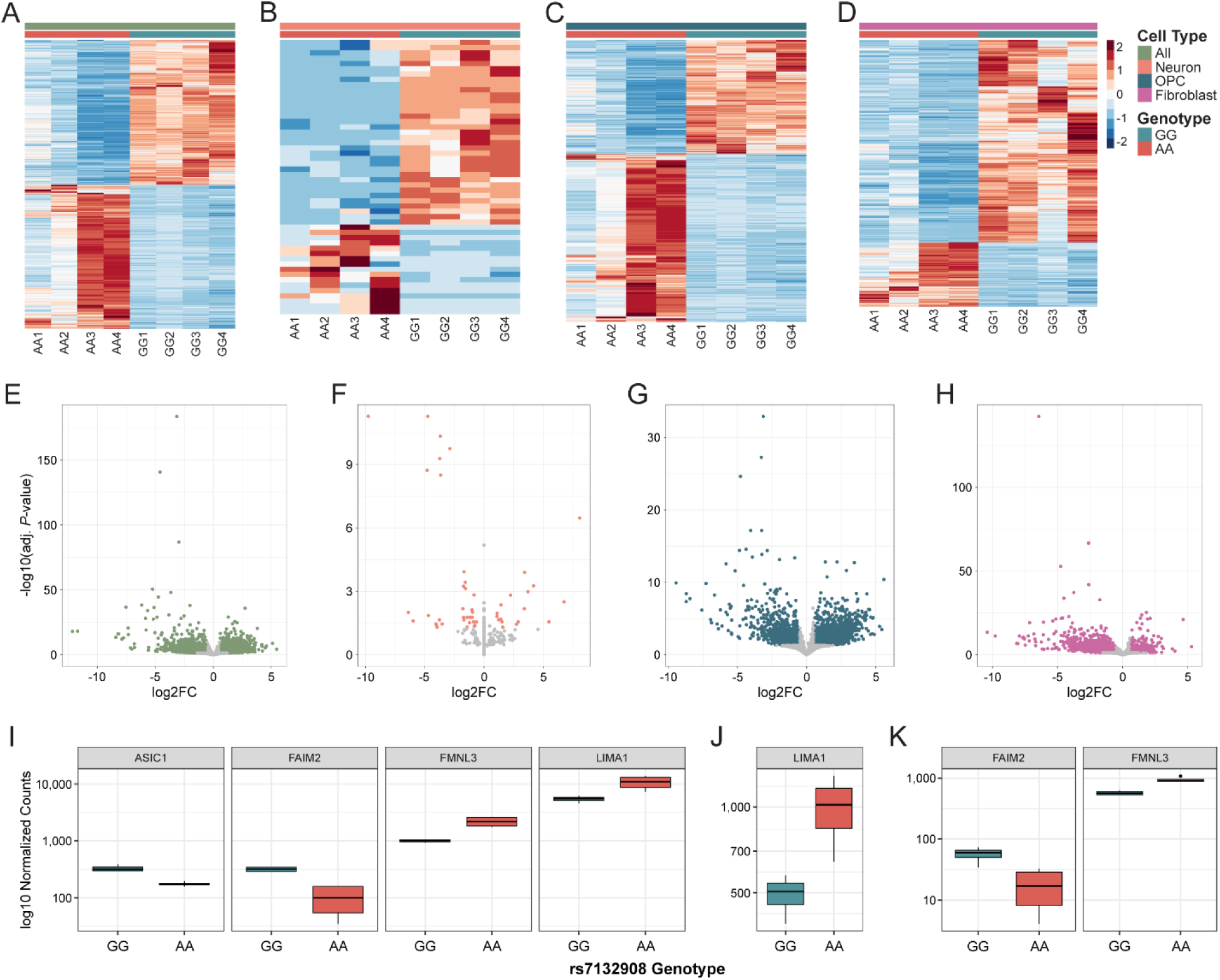
rs7132908 genotype influences gene expression in ESC-derived hypothalamic cell types. A-D) Heatmaps depicting significantly differentially expressed genes (adjusted P-value < 0.05, |log2 fold change| > 0.58) due to the rs7132908 risk allele in all cells (A), neurons (B), OPCs (C), and fibroblasts (D) produced by ESC differentiation. E-H) Volcano plots of adjusted P-values (-log10) and fold change (log2) of genes tested for differential expression due to the rs7132908 obesity risk allele in all cells (E), neurons (F), OPCs (G), and fibroblasts (H) produced by ESC differentiation. Colored dots indicate significantly differentially expressed genes (adjusted P-value < 0.05, |log2 fold change| > 0.58) due to the rs7132908 risk allele and grey dots indicate genes with no significant differences in expression. I-K) Box plots of gene expression (log10 normalized counts) for genes in the rs7132908 TAD that were significantly differentially expressed (adjusted P-value < 0.05, |log2 fold change| > 0.58) in all cells (I), OPCs (J), and fibroblasts (K). See also Figure S3 and S4.

**Supplemental Figure 3.**
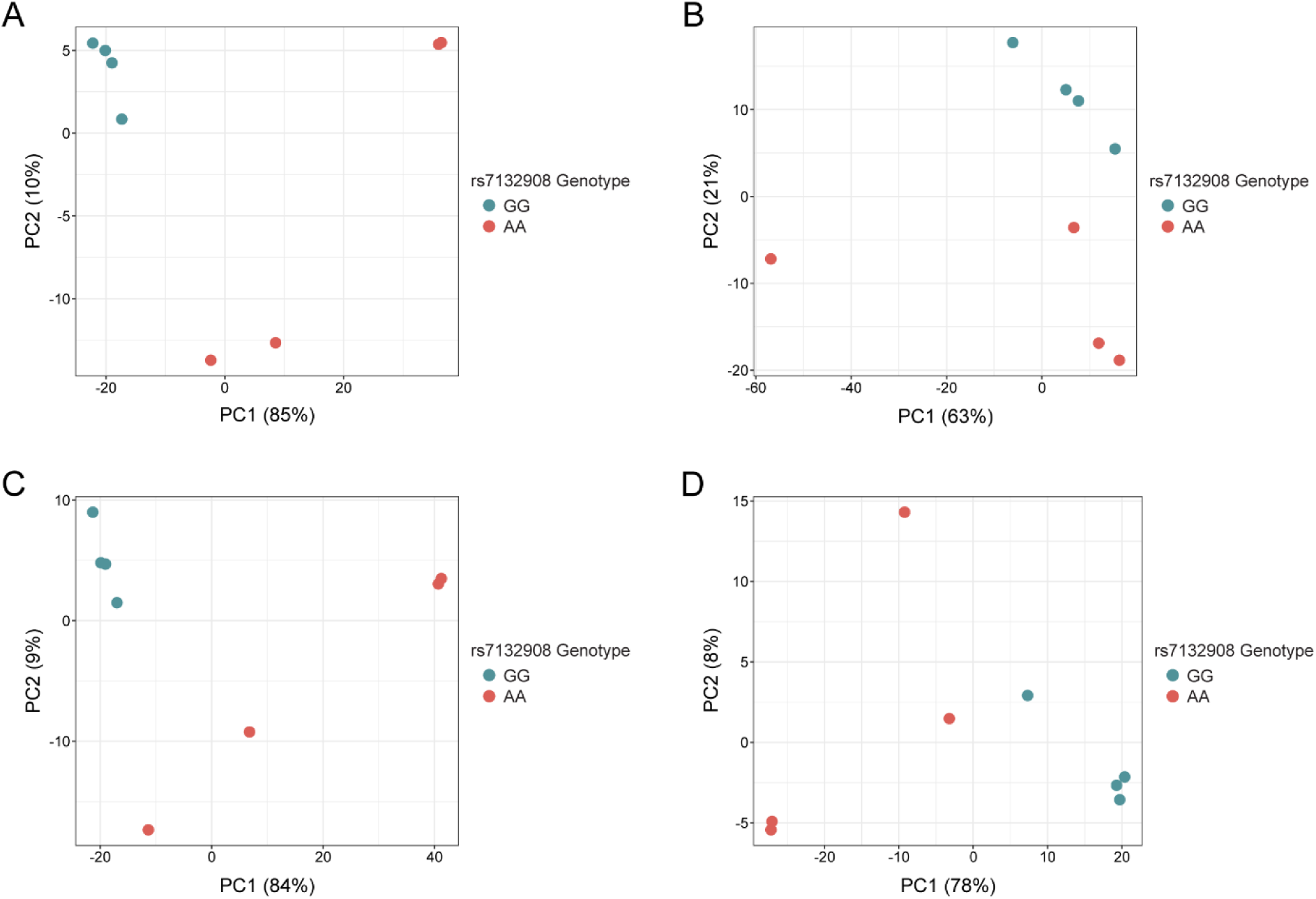
Hypothalamic single-nucleus RNA-seq differential expression analysis, related to Figure 7. **A-D)** PCA plots of single-nucleus RNA-seq libraries (GG n = 4, AA n = 4) when considering all cells (A), neurons (B), OPCs (C), and fibroblasts (D).

Next, we identified genes differentially expressed within each annotated cell type. rs7132908 genotype explained 21%, 84%, and 78% of the variation between replicate samples in the neurons, OPCs, and fibroblasts, respectively (**Fig. S3B-D**). In neurons, 52 genes were differentially expressed, with 35 down-regulated in neurons homozygous for the risk allele and 17 up-regulated (**Fig. 7B**, **Fig. 7F, Supp. Table 8**). In OPCs, 2,678 genes were differentially expressed, with 1,084 down-regulated in OPCs homozygous for the risk allele and 1,594 up-regulated (**Fig. 7C**, **Fig. 7G, Supp. Table 8**), while in fibroblasts, 1,911 genes were differentially expressed, with 1,450 down-regulated in fibroblasts homozygous for the risk allele and 461 up-regulated (**Fig. 7D**, **Fig. 7H, Supp. Table 8**). When considering genes located in the same TAD as rs7132908^51^, no genes were differentially expressed in neurons, while one gene was differentially expressed in OPCs (*LIMA1* up-regulated) (**Fig. 7J**), and two genes were differentially expressed in fibroblasts (*FAIM2* down-regulated; *FMNL3* up-regulated) (**Fig. 7K**). Functional enrichment analyses of up-regulated genes in both OPCs and fibroblasts identified similar Gene Ontology terms^52,53^, including the biological processes of cell death and apoptosis (**Supp. Table 9**), while processes such as nervous system development, neuron differentiation, and neuron projection development were enriched among down-regulated genes (**Supp. Table 9**). However, the comparably shorter lists of significantly up- and down-regulated genes in neurons did not identify any significantly enriched biological processes.

As our sequencing efforts only captured transcriptional differences at three timepoints, we were therefore motivated to quantify *FAIM2* expression in all cells throughout the 40-day hypothalamic neuron differentiation using quantitative real-time polymerase chain reaction (RT-qPCR). *FAIM2* expression peaked around day 14 in cells homozygous for either rs7132908 allele (**Fig. S4A-B**), which represents the hypothalamic neural progenitor phase of the differentiation (**Fig. 3A**). We also characterized *FAIM2* expression *in vivo* using our primary human pediatric (donor ages 4-14 years old, average age = 7.5) hypothalamus tissue RNA-seq data and determined that *FAIM2* was highly expressed (median TPM = 415.66, n = 4) (**Fig. S4C, Supp. Table 10**).

**Supplemental Figure 4.**
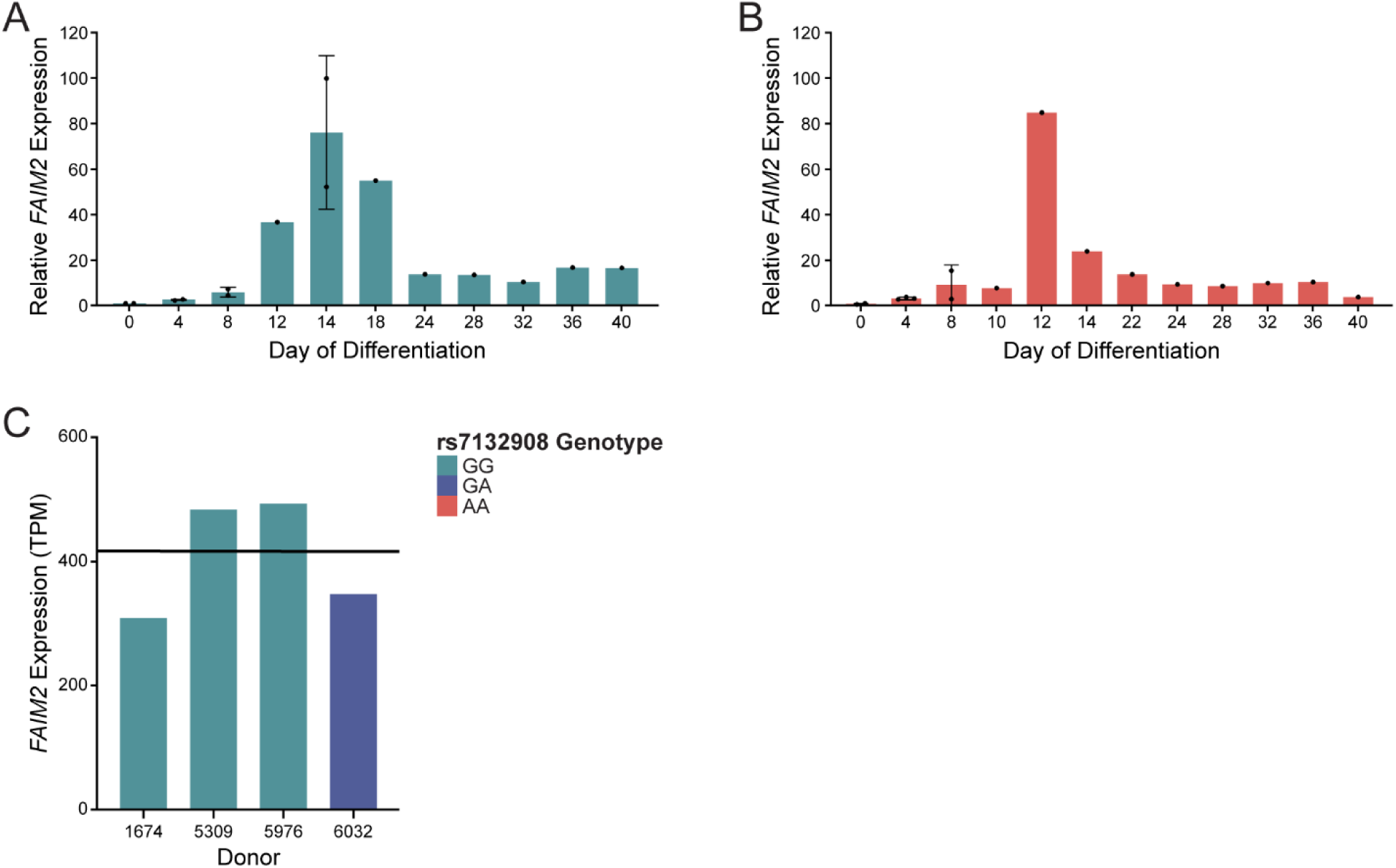
FAIM2 expression, related to Figure 7. **A-B)** Relative normalized FAIM2 mRNA expression in cells homozygous for the rs7132908 non-risk allele (A) and obesity risk allele (B) measured by RT-qPCR throughout ESC differentiation to hypothalamic neurons. FAIM2 expression was normalized to 18S ribosomal RNA expression. Relative FAIM2 expression was calculated relative to non-risk allele cells on day 0. Data are represented as mean ± SD when n > 1. C) FAIM2 expression (TPM) in primary human pediatric hypothalamus tissue. Black horizontal line indicates median expression (n=4). Blue bars indicate donors homozygous for the rs7132908 non-risk G allele and the indigo bar indicates a donor heterozygous at rs7132908.

## DISCUSSION

The chr12q13 locus was first associated with variation in adult BMI and weight in 2009^65^, BMI as a longitudinal trait during childhood (ages 3-17) in 2012^30^, and childhood obesity as a dichotomous trait by our meta-analysis with the EGG consortium in 2012^22^. It was also noted that the genotypic risk effect at the chr12q13 locus during childhood decreased as age increased^30^, which suggests this locus may regulate age-dependent pathways in early childhood and could explain why this locus is more pronounced in childhood than adulthood. More than 1,000 independent loci are now associated with measurements of obesity^25^ and only a few have been studied extensively enough to pinpoint a causal variant and implicate effector genes, such as the *FTO*^26–28^ and *2q24.3* loci^66^.

A recent global functional investigation of BMI-associated SNPs in 3’ UTRs included the *FAIM2* 3’ UTR variant rs7132908 in their study and found that the obesity-associated risk A allele disrupted miRNA binding activity of miR-330-5p in hamster ovary cells and human subcutaneous preadipocytes, leading to an increase in *FAIM2* expression^67^. These results may however not accurately reflect regulation of *FAIM2* expression *in vivo* as this gene is primarily expressed in the brain; furthermore, this miRNA product is a passenger strand that is typically found in lower abundance due to degradation after its complementary guide miRNA is loaded into an Argonaute protein during miRNA processing^68^. More recently, others carrying out global analyses have implicated an enhancer in the region harboring rs7132908 with a luciferase reporter assay and found that, in mouse neuronal hypothalamus cells, the obesity risk A allele significantly decreased enhancer activity with a minimal promoter^69^, consistent with our results.

We first utilized a luciferase reporter assay in a model tolerant of transfection, human primary astrocytes, to characterize the regulatory effects of the non-coding region surrounding rs7132908 on genes within its TAD that were nominated by our variant-to-gene mapping efforts in this cell type. Ectopic expression of this non-coding region with the rs7132908 non-risk G allele significantly increased reporter expression driven by the *FAIM2* promoter, while the obesity risk A allele significantly decreased reporter expression driven by the *FAIM2* promoter. These results further support *FAIM2* as the lead candidate effector gene at the chr12q13 obesity locus and rule out *LIMA1* and *RACGAP1* as effector genes specifically in astrocytes. While this tractable *in vitro* model was used to characterize *cis*-regulatory activity, we note that BMI-associated SNPs are not enriched in astrocyte-specific *cis*-regulatory elements and that FAIM2 likely functions primarily in neurons^37–41^.

Next, we used a human stem cell-derived model of hypothalamic neural progenitors and neurons. Given the dynamic changes in genomic architecture that occur during stem cell differentiation, we characterized gene expression and chromatin accessibility at several timepoints. We used bulk sequencing approaches when the cells were expected to be homogenous and leveraged multi-omic single-nucleus RNA-seq and ATAC-seq at the post-differentiation hypothalamic neuron stage to capture cell type-specific differences in this now heterogeneous model. We determined that the rs7132908 obesity risk A allele led to significant differential expression of 0 TAD genes in ESCs, 5 TAD genes in hypothalamic neural progenitors (*AQP2, COX14, FAIM2, LARP4,* and *TMBIM6*), 1 TAD gene in OPCs (*LIMA1*), and 2 TAD genes in fibroblasts (*FAIM2* and *FMNL3*). These results, in combination with our observation that rs7132908 is not accessible in ESCs, suggest that rs7132908 does not regulate gene expression in stem cells. These results also implicate different effector genes depending on cell type, in agreement with the luciferase assay results where enhancer activity was observed for *FAIM2* in primary astrocytes but not in HEK293Ts. Only *FAIM2* was implicated in more than one cell type and its expression was consistently down-regulated with the obesity risk A allele. Taken together, we demonstrated that rs7132908 resides within a *cis­*-regulatory element that confers allele-specific and cell type-specific effects on the expression of *FAIM2* and other genes within its TAD.

We did not observe large differences in accessibility at rs7132908 due to genotype in any cell type. Therefore, significant changes in effector gene expression are likely due to differences in transcription factor binding affinity. We predicted that the rs7132908 risk A allele disrupts binding of 12 transcription factors, many of which are known to be both activators and repressors and are ubiquitously expressed. Further investigation is warranted to determine which specific transcription factors bind at and temporally regulate gene expression at the chr12q13 locus.

In this study, we made the striking observation that the rs7132908 obesity risk A allele decreased the proportion of hypothalamic neurons produced by stem cell differentiation. In addition, we observed that the obesity risk A allele led to up-regulation of cell death and apoptosis gene sets and down-regulation of neuron development gene sets. However, we did not observe that any orexigenic or anorexigenic neuronal cell cluster or subpopulation was more severely decreased, highlighting the need for more experiments to determine how the rs7132908 obesity risk allele could increase appetite and risk of childhood obesity.

Our working hypothesis is that rs7132908 regulates *FAIM2* and possibly other genes that are required for normal anorexigenic neuron development or survival at a crucial timepoint in development. When *FAIM2* expression was highest in hypothalamic neural progenitors, its expression was approximately 50% less in progenitors homozygous for the rs7132908 obesity risk allele. FAIM2 protects neurons from Fas-induced apoptosis^37,38^ and regulates neurite outgrowth^39^, neuroplasticity^40^, and synapse formation^41^. While *Faim2* null mice have only been previously used to study neurological^70–73^ and immune^74^ phenotypes, one study reported that *Faim2* null mice at 10-12 weeks of age and fed a standard diet *ad libitum* did display subtle increases in fat content^70^. Rodent studies have also demonstrated that *Faim2* expression increased in the hypothalamic arcuate nucleus in response to restricted food intake^75^ and food deprivation^76^. Future work must be dedicated to directly test our hypothesis that *FAIM2* is a causal effector gene for childhood obesity and a more complex model system may be appropriate for such studies.

There are several limitations to our study to consider. First, although our ESC-derived *in vitro* model of hypothalamic neurogenesis expresses some appropriate marker genes, it likely does not fully recapitulate the hypothalamus during childhood. We also generated non-neuronal cell types (OPCs and fibroblasts) that correlated most highly with cultured fibroblasts in the GTEx RNA-seq database^36^ but still expressed neuronal markers (*MAP2* and *TUBB3*) at some level, likely due to exposure to neuron maturation cell culture medium for 26 days. While we reported changes in gene expression and chromatin accessibility in these additional cell types, they may not be as biologically relevant. Second, we used the female H9 ESC line which prevented us from detecting sex-specific differences. Third, we did not investigate the effects of the rs7132908 obesity risk A allele *in vivo*. We were able to obtain four pediatric hypothalamus tissue samples, but with just three homozygous for the rs7132908 non-risk allele and only one heterozygote, this sample size was insufficient for allele-specific expression or eQTL analyses. In the future, increased accessibility to human pediatric hypothalamus tissue would aid investigation at the chr12q13 childhood obesity locus.

Overall, we functionally validated rs7132908 as a causal SNP at one of the strongest but commonly overlooked childhood obesity GWAS loci, implicated *FAIM2* and other cell type-specific effector genes, and nominated pathways acting downstream of the SNP involving nervous system development and cell death. We have also generated datasets from primary astrocytes and multiple timepoints throughout hypothalamic neuron differentiation, including multi-omic single-nucleus RNA-seq and ATAC-seq data, that will serve as a resource to aid investigation of other loci and traits relevant to our cell models. This progress towards characterizing the precise mechanism underlying the association between the chr12q13 genomic region and obesity should in turn enable future work with this key locus and guide comparable efforts to validate other causal genes and to ultimately identify therapeutic targets.

## ACKNOWLEDGEMENTS

We thank Dr. Bill Manley, Madeleine Salvatore, and Danny Frederick for training in stem cell culture; Dr. Guo-li Ming and Dr. Sarshan Pather for sharing annotated single-cell human hypothalamus transcriptome data and code; Dr. Dhruv Sareen and Andrew Gross for sharing updated differentiation protocols and providing technical support; Dr. Daniel Beiting for training in RNA-seq analysis; Gina Pacella for providing experiment design guidance; Dr. Jonathan Schug, Dr. Winter Bruner, and Mitchell Conery for providing bioinformatic analysis guidance; Dr. Alexis Crockett for providing mouse brain tissue for a DNA and RNA extraction pilot study; Dr. Shaon Sengupta for allowing use of their benchtop homogenizer; Kenyaita Hodge for preparing ATAC-seq libraries; Sumei Lu for preparing RNA-seq libraries; Children’s Hospital of Philadelphia Human Pluripotent Stem Cell Core for CRISPR and karyotyping services; Children’s Hospital of Philadelphia Center for Applied Genomics for genotyping, CNV analysis, and single-nucleus sequencing services; Children’s Hospital of Philadelphia Flow Cytometry Core for cell sorting services; University of Pennsylvania Genomic and Sequencing Core DNA Sequencing Laboratory for Sanger sequencing services; and University of Pennsylvania Cell Center Services for stockroom services. Human tissue was obtained from the NIH Neurobiobank at the University of Maryland, Baltimore, MD. Some figures were created with BioRender.com. S.H.L is supported by the NICHD (F31 HD105404). S.F.A.G. is supported by the NICHD (R01 HD056465), the NIDDK (UM1 DK126194), and the Daniel B. Burke Endowed Chair for Diabetes Research.

## AUTHOR CONTRIBUTIONS

S.H.L. and S.F.A.G. conceived the project. S.H.L. designed the experiments. S.H.L., C.M.V., N.D., and K.C. performed cell culture. S.H.L. and C.M.V. collected relevant cell materials. S.H.L. processed frozen human tissue. S.H.L. and C.M.V. conducted cell line validation. S.H.L. optimized human primary astrocyte transfection. S.H.L., C.M.V., and N.D. performed luciferase assays. S.H.L. performed luciferase assay data analysis. J.A.M. designed and performed CRISPR. S.H.L. and K.C. performed immunocytochemistry and imaging. S.H.L. and K.B. prepared bulk RNA-seq libraries. J.A.P. prepared bulk ATAC-seq libraries. J.A.P. and S.H.L. prepared Hi-C libraries. S.H.L. prepared nuclei for single-nucleus RNA-seq and ATAC-seq. J.A.P. sequenced bulk RNA-seq, ATAC-seq, and Hi-C libraries. S.H.L., K.B.T, and A.C. performed bulk RNA-seq analyses. M.C.P., K.B.T., and A.C. performed bulk ATAC-seq analyses. K.B.T. performed transcription factor, colocalization, and Hi-C analyses. S.H.L., K.B.T., and M.A.W. performed single-nucleus RNA-seq analyses. K.B.T. and S.H.L. performed single-nucleus ATAC-seq analyses. S.H.L. performed RT-qPCR and data analysis. S.F.A.G., M.C.P., A.D.W., S.A.A., and J.A.P. provided critical feedback and supervision. S.H.L. and S.F.A.G. wrote the original manuscript draft. All authors reviewed and edited the final manuscript.

## STAR METHODS

### Resource availability

#### Lead contact

Further information and requests for resources and reagents should be directed to and will be fulfilled by the lead contact, Struan F. A. Grant.

#### Materials availability

Vectors (pGL4.10[*luc2*]-rs7132908G-*FAIM2*, pGL4.10[*luc2*]-rs7132908A-*FAIM2*, pGL4.10[*luc2*]-*FAIM2*, pGL4.10[*luc2*]-rs7132908G-*LIMA1*, pGL4.10[*luc2*]-rs7132908A-*LIMA1*, pGL4.10[*luc2*]-*LIMA1*, pGL4.10[*luc2*]-rs7132908G-*RACGAP1*, pGL4.10[*luc2*]-rs7132908A-*RACGAP1*, pGL4.10[*luc2*]-*RACGAP1*, and gRNA_Cloning-rs7132908gRNA) and cell lines (WA09 (H9) rs7132908 AA human embryonic stem cell clones 2.1, 9.1, and 10.1) generated in this study will be available from the lead contact with a completed Materials Transfer Agreement. This study did not generate any other new unique reagents.

#### Data and code availability

Genotyping, Hi-C, RNA-seq, ATAC-seq, single-nucleus RNA-seq, and single-nucleus ATAC-seq data have been deposited at Gene Expression Omnibus (GEO) and are publicly available as of the date of publication. Accession numbers are listed in the key resources table. This paper does not report original code. Any additional information required to reanalyze the data reported in this paper is available from the lead contact upon request.

### Experimental model and subject details

#### Primary astrocyte model

Primary Normal Human Astrocytes (NHA) of unknown sex were obtained from Lonza as cryopreserved cells. The cells were obtained at passage 1 and used before passage 10, as recommended. They were cultured following Lonza technical instructions in Lonza Astrocyte Growth Medium and in a humidified incubator at 37°C with 5% CO_2_. For thawing, cells were thawed quickly at 37°C, resuspended, and added slowly to an excess of warmed medium to seed at approximately 6,500 cells/cm^2^ in a T75 flask. For passaging, 70-80% confluent cells were washed with 30 mM HEPES buffered saline solution in water, incubated at 37°C with 0.025% trypsin-EDTA in DPBS for 3-4 minutes or until 90% of the cells rounded up, treated with 2 volumes of 5% FBS in DPBS to neutralize the trypsin, rinsed off the culture vessel with gentle pipetting, pelleted by centrifugation at 160 rcf for 5 minutes at 4°C, and then resuspended and seeded at the desired density. The cells were cultured in T75 flasks, 6-well plates, and 24-well plates. For freezing, cells were lifted as for passaging, resuspended to 1,000,000 cells/mL in FBS with 10% DMSO, frozen in 1 mL aliquots at -1°C/minute, and stored long-term in liquid nitrogen. The cells tested negative for mycoplasma contamination (**Fig. S5D**).

**Supplemental Figure 5.**
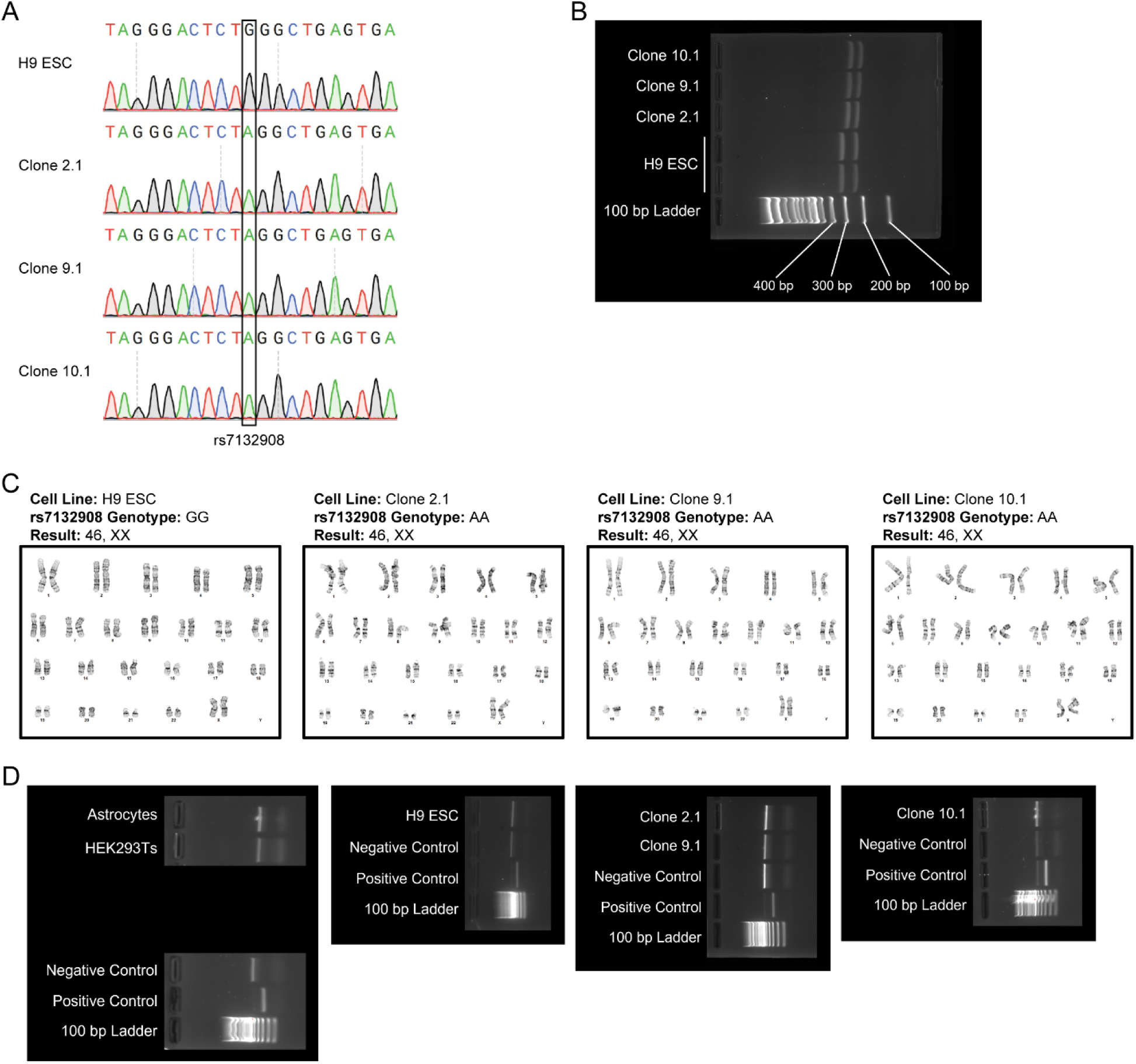
Validation of experimental models, related to STAR Methods. A) Electropherograms produced by Sanger sequencing around rs7132908 in ESC lines. B) BfaI restriction enzyme digestion screening in ESC lines. H9 ESCs have one BfaI restriction site in the PCR product around rs7132908, where digestion should produce two bands of 320 bp and 248 bp. After CRISPR to introduce the rs7132908 obesity risk A allele, a second BfaI restriction site is introduced, where digestion should produce three bands of 294 bp, 248 bp, and 26 bp (not pictured). C) G-band karyotyping reports for ESC lines. D) Mycoplasma PCR detection results for all experimental models. Cell lines with bands matching the size of the negative control are not contaminated with mycoplasma.

#### HEK293T model

293T human female cells were obtained from ATCC as cryopreserved cells (ATCC Cat# CRL-3216; RRID: CVCL_0063). They were cultured following ATCC product information in Dulbecco’s Modified Eagle’s Medium (DMEM) with 10% FBS, 1X Antibiotic-Antimycotic, and 2 mM L-glutamine and in a humidified incubator at 37°C with 5% CO_2_. For thawing, cells were thawed quickly at 37°C, resuspended, added slowly to an excess of warmed medium, pelleted by centrifugation at 125 rcf for 7 minutes at 25°C, resuspended in warmed medium, and seeded at approximately 17,500 cells/cm^2^ in a 10 cm dish. For passaging, 90% confluent cells were washed with PBS, incubated at 37°C with 0.25% trypsin-EDTA for 4-5 minutes, treated with 2 volumes of medium to neutralize the trypsin, pelleted by centrifugation at 1,200 rcf for 2 minutes at 25°C, and then resuspended and seeded at the desired density. The cells were cultured in 10 cm dishes, 6-well plates, and 24-well plates. For freezing, cells were lifted as for passaging, resuspended to 1,000,000 cells/mL in medium with 5% DMSO, frozen in 1 mL aliquots at -1°C/minute, and stored long-term in liquid nitrogen. The cells tested negative for mycoplasma contamination (**Fig. S5D**).

#### ESC model

WA09 (H9) human female embryonic stem cells were obtained from the WiCell Research Institute as cryopreserved cells (WiCell Lot# DL-05; RRID: CVCL_9773). Before use, the cells were authenticated with short tandem repeat analysis to confirm cell line identity. They were cultured following WiCell protocols in mTeSR1 medium, on Matrigel hESC-qualified matrix, and in a humidified incubator at 37°C with 5% CO_2_. During CRISPR editing, the cells were briefly cultured on Matrigel Growth Factor Reduced Basement Membrane Matrix diluted in IMDM and mouse embryonic fibroblasts (MEFs) and in DMEM/F12 medium supplemented with 15% volume KnockOut Serum Replacement, 100 µM non-essential amino acids, 1 mM sodium pyruvate, 2 mM L-glutamine, 50 U/mL penicillin-streptomycin, 0.1 mM β-mercaptoethanol, and 10 ng/mL human bFGF. For thawing, cells were thawed quickly at 37°C, resuspended, added slowly to an excess of warmed medium, pelleted by centrifugation at 200 rcf for 5 minutes at 25°C, resuspended in warmed medium, and seeded into 1 well of a 6-well plate. For passaging as colonies, cells in large colonies were washed with Versene, incubated at room temperature with Versene for 6-9 minutes, rinsed off the culture vessel with medium and gentle pipetting, and then split across new culture vessels, generally using a 1:12 ratio. For passaging as single cells, cells in large colonies were washed with DPBS, incubated at 37°C with Accutase for 2-5 minutes, treated with 2 volumes of medium to neutralize the Accutase, pelleted by centrifugation at 200 rcf for 4 minutes at 25°C, and then resuspended and seeded at the desired density. For passaging when cultured on MEFs, MEFs were removed by incubating with TrypLE Express Enzyme for 3 minutes at room temperature. 10 µM ROCK Inhibitor Y-27632 was added to the medium for 24 hours after thawing or passaging as single cells. The cells were cultured in 10 cm dishes, T25 flasks, 6-well plates, and 24-well plates. For freezing, cells were lifted as colonies as for passaging, pelleted by centrifugation at 200 rcf for 4 minutes at 25°C, resuspended in 2 mL mFreSR medium/lifted well of a 6-well plate, frozen in 1 mL aliquots at -1°C/minute, and stored long-term in liquid nitrogen. The cells were validated with karyotyping (**Fig. S5C**) and tested negative for mycoplasma contamination (**Fig. S5D**).

#### Pediatric post-mortem brain tissue

Frozen human pediatric hypothalamus tissue from 4 post-mortem individuals were obtained. The tissue donors included a 4-year-old male, 8-year-old male, 4-year-old female, and 14-year-old female, all classified as white and with no clinical diagnoses. The number of samples was limited by tissue availability.

### Method details

#### Mycoplasma contamination testing

Cells were cultured in the absence of antibiotics for several days and until 90-100% confluent. Medium was then collected and used to detect mycoplasma by PCR using the LookOut Mycoplasma PCR Detection kit with JumpStart Taq DNA polymerase, following manufacturer’s instructions. PCR products, including positive and negative controls, were visualized with gel electrophoresis. Band sizes from experimental samples were compared to the negative control to determine that all cell cultures were negative for mycoplasma contamination (**Fig. S5D**).

#### Bulk ATAC-seq library preparation

ATAC-seq libraries were prepared from primary astrocytes with 3 technical replicates, the rs7132908 non-risk G allele ESCs with 3 technical replicates and the rs7132908 risk A allele ESCs with 3 biological replicates. 50,000-100,000 cells from each replicate were centrifuged at 550 rcf for 5 minutes at 4°C to pellet. Each cell pellet was washed with cold PBS and resuspended in 50 μL cold lysis buffer (10 mM Tris-HCl, pH 7.4, 10 mM NaCl, 3 mM MgCl_2_, and 0.1% IGEPAL CA-630) then immediately centrifuged at 550 rcf for 10 minutes at 4°C. Nuclei were resuspended in transposition reaction mix (25 μL 2X Tagment DNA Buffer, 2.5 μL TDE1 Tagment DNA Enzyme, and 22.5 μL nuclease-free water) on ice, then incubated for 45 minutes at 37°C. The tagmented DNA was then purified using the Qiagen MinElute PCR Purification kit and eluted in 10.5 μL elution buffer. 10 μL of each purified tagmented DNA sample was amplified with PCR using the Nextera DNA CD Indexes kit and NEBNext High-Fidelity PCR Master Mix for 12 cycles to generate each library. The libraries were purified using AMPure XP beads at a 1.8X concentration. Library concentrations were measured with Qubit dsDNA High Sensitivity Assays. The completed libraries were assessed with the Agilent Bioanalyzer DNA 1000 kit and 2100 Bioanalyzer Expert software (RRID: SCR_019715). Completed libraries were pooled and sequenced on the Illumina NovaSeq 6000 platform using paired-end 51 bp reads.

#### Hi-C library preparation

Hi-C libraries were prepared from primary astrocytes with two technical replicates using the Arima-HiC kit, following manufacturer’s instructions and as previously described^47^. In brief, cells were crosslinked with formaldehyde and then chromatin was digested with multiple restriction enzymes. The purified proximally-ligated DNA was then sheared and 200-600 bp DNA fragments were selected with AMPure XP beads. The size-selected fragments were then enriched using Enrichment Beads and then converted to Illumina-compatible sequencing libraries using the Swift Accel-NGS 2S Plus DNA Library kit and Swift 2S Indexing kit. The libraries were assessed using the Agilent Bioanalyzer DNA 1000 kit and 2100 Bioanalyzer Expert software (RRID: SCR_019715) and the KAPA Library Quantification kit. Completed libraries were pooled and sequenced on the Illumina NovaSeq 6000 platform using paired-end 101 bp reads.

#### RNA extraction from cells

To extract RNA from cultured cells for RNA-seq or RT-qPCR, cells were lifted and resuspended in TRIzol. RNA was extracted from each TRIzol sample with the Zymo Direct-zol RNA Miniprep kit, following manufacturer’s instructions, with recommended DNase I treatment.

#### DNA and RNA extraction from tissue

DNA and RNA were extracted from frozen human pediatric hypothalamus tissue samples in parallel. Each tissue sample was homogenized in DNA/RNA Shield in 2 mm ZR BashingBead Lysis Tubes with a FastPrep-24 5G high-speed benchtop homogenizer at 10 m/s at room temperature for 45 seconds. DNA and RNA were then extracted using the Zymo Quick-DNA/RNA Miniprep Plus kit, following manufacturer’s instructions.

#### Bulk RNA-seq library preparation

RNA extracted from each cell line and tissue sample was quantified and assessed with the Agilent Bioanalyzer RNA 6000 Nano kit and 2100 Bioanalyzer Expert software (RRID: SCR_019715). Cell line samples with an RNA integrity number (RIN) greater than 7 and tissue samples with a RIN greater than 5 were used for RNA-seq library preparation. RNA-seq libraries were prepared from each tissue sample with 3 technical replicates, primary astrocytes with 3 technical replicates, the rs7132908 non-risk G allele ESCs with 2 technical replicates, the rs7132908 risk A allele ESCs with 3 biological replicates, and hypothalamic neural progenitors with either allele from two independent differentiations (biological replicates) with 3 technical replicates. 40 ng to 1 µg of each RNA sample was used as input, depending on RNA extraction yield. Ribosomal RNA was depleted using the QIAseq FastSelect RNA Removal kit, following manufacturer’s instructions. Libraries were prepared using the NEBNext Ultra II Directional RNA Library Prep for Illumina kit, NEBNext Oligos for Illumina (Dual Index Primers Set 1), and AMPure XP beads, following manufacturer’s instructions. Library concentrations were quantified with Qubit dsDNA High Sensitivity Assays. 5 ng of each library was used for assessment with the Agilent Bioanalyzer DNA 1000 kit and 2100 Bioanalyzer Expert software (RRID: SCR_019715). If the electropherogram did not display a narrow sample distribution around 300 bp, an additional bead cleanup or column purification was used to remove any contaminating primers, adapter-dimers, or large fragments generated by over-amplification. Completed libraries were pooled and sequenced on the Illumina NovaSeq 6000 platform using paired-end 51 bp reads.

#### Primary astrocyte transfection optimization

To optimize transfection of the primary astrocytes, we transfected with varying amounts of LentiCRISPRv2-mCherry vector DNA, which was a gift from Agata Smogorzewska (Addgene Cat# 99154; http://n2t.net/addgene:99154; RRID: Addgene_99154), Lipofectamine LTX, and PLUS Reagent and then quantified transfection efficiency and cell viability with flow cytometry in two separate experiments. Primary astrocytes were seeded at 50,000 cells/well in a 24-well plate and maintained until they reached 70-80% confluence. Lipofectamine LTX-DNA complexes with PLUS Reagent were prepared following manufacturer’s instructions in Opti-MEM so that each well would receive either 0 ng, 250 ng, 500 ng, or 750 ng vector DNA, 1 µL PLUS Reagent/1 µg of vector DNA, and either a 1:1, 1:2, 1:2.5, 1:3, 1:4, or 1:5 vector DNA (µg):Lipofectamine LTX (µL) ratio.

Approximately 22 hours post-transfection, the cells were lifted, resuspended in PBS, fixed in 2% paraformaldehyde for 10 minutes at room temperature, resuspended in PBS, strained using a 35 μm strainer, and then counted using a CytoFLEX S N2-V3-B5-R3 Flow Cytometer. 10,000 events were collected for each condition and gating was set using the non-transfected control condition. Percent single cell events was calculated by dividing the number of single cell events by all events (10,000). Percent cell viability was then calculated by dividing the percent single cell events for each condition by the average percent single cell events for 2 replicates of non-transfected controls. Transfection efficiency was calculated by dividing the number of mCherry+ single cell events by the number of single cell events in each condition. This optimization experiment determined that ideal conditions for transfecting primary astrocytes at 70-80% confluence in a 24-well plate for 22 hours are 750 ng vector DNA, 0.75 µL PLUS Reagent, and 1.875 µL Lipofectamine LTX (1:2.5 ratio) diluted in Opti-MEM for a total volume of 50 µL/well, which was used for all future primary astrocyte transfection experiments. These transfection conditions yielded high transfection efficiency (11.26%) when considering that the expected efficiency is 5-12%^77^ and high cell viability (85.69%) (**Fig. S6A-B**).

**Supplemental Figure 6.**
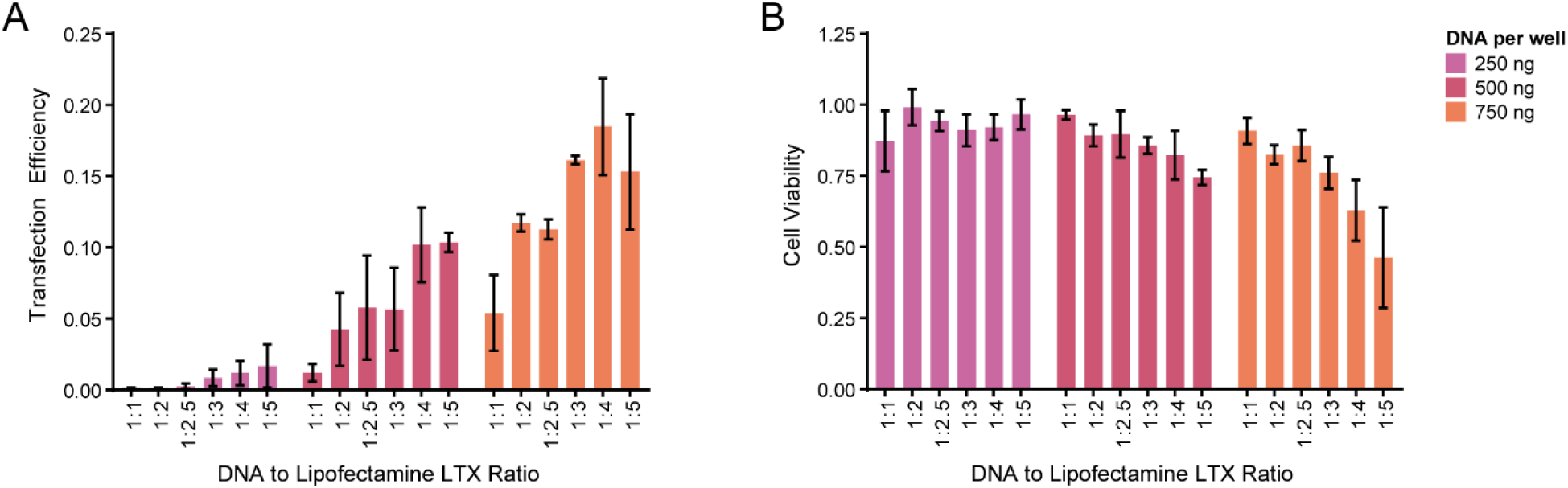
Primary astrocyte transfection optimization, related to STAR Methods. A) Transfection efficiency resulting from transfecting with 250, 500, or 700 ng DNA per well and varying DNA to Lipofectamine LTX ratios (n = 2 biological replicates). B) Cell viability resulting from transfecting with 250, 500, or 700 ng DNA per well and varying DNA to Lipofectamine LTX ratios (n = 2 biological replicates). Data are represented as mean ± SD.

#### Generation of luciferase assay vectors

The ENCODE consortium’s ‘Registry of candidate *cis-*Regulatory Elements’ (version 1) (RRID: SCR_006793) annotated a cell type-agnostic regulatory element with a distal enhancer-like signature surrounding rs7132908 at chr12:50,262,620-50,263,581 (GRCh37)^50^. To generate a DNA fragment containing this sequence with an additional 50 bp flanking each side for cloning, we designed PCR primers to amplify this region of interest and used a *FAIM2* 3’ UTR miRNA target clone (purchased from GeneCopoeia) as the PCR template and NEBNext High-Fidelity PCR Master Mix. To generate DNA fragments containing the *FAIM2*, *LIMA1*, and *RACGAP1* promoter sequences, we also designed PCR primers to amplify these regions and used promoter clones (purchased from GeneCopoeia) as PCR templates. The promoterless pGL4.10[*luc2*] firefly luciferase reporter vector (purchased from Promega) was linearized at the multiple cloning site upstream of the *luc2* reporter gene using the XhoI restriction enzyme. Each PCR product and the linearized plasmid were extracted after visualization with gel electrophoresis with the NEB Monarch DNA Gel Extraction kit to ensure that a fragment of correct length was purified. The putative enhancer region containing rs7132908 and each promoter were inserted at the multiple cloning site of pGL4.10[*luc2*] using the Codex Gibson Assembly HiFi HC 1-Step kit to generate pGL4.10[*luc2*]-rs7132908G-*FAIM2*, pGL4.10[*luc2*]-rs7132908G-*LIMA1*, and pGL4.10[*luc2*]-rs7132908G-*RACGAP1* vectors. Each promoter alone was also inserted at the multiple cloning site to generate pGL4.10[*luc2*]-*FAIM2,* pGL4.10[*luc2*]-*LIMA1,* and pGL4.10[*luc2*]-*RACGAP1* control vectors. Each Gibson Assembly product was used to transform NEB Stable Competent *E. coli* which were then plated on LB agarose plates with 100 µg/mL ampicillin to select for successfully transformed colonies. Bacterial plates were incubated overnight at 37°C and then individual colonies were selected for overnight growth in LB broth with 100 µg/mL ampicillin at 30°C with shaking at 250 rpm. Vector DNA was extracted from each overnight culture using the Qiagen QIAprep Spin Miniprep kit and then Sanger sequenced on both strands throughout the modified region to confirm successful insertion and sequence. Electropherograms and sequence files produced from Sanger sequencing were analyzed using SnapGene software (RRID: SCR_015052). Once vectors with perfect sequences were identified, we used the NEB Q5 Site-Directed Mutagenesis kit to introduce the childhood obesity risk A allele at rs7132908 and generate pGL4.10[*luc2*]-rs7132908A-*FAIM2*, pGL4.10[*luc2*]-rs7132908A-*LIMA1*, and pGL4.10[*luc2*]-rs7132908A-*RACGAP1* vectors. We used Sanger sequencing on both strands throughout the modified region to confirm successful mutagenesis and lack of polymerase errors. Bacteria glycerol stocks were prepared to store each transformed strain with verified sequences long-term. Each experimental vector, the unmodified pGL4.10[*luc2*] control vector, and pRL-TK (purchased from Promega) co-transfection control vector were then purified for transfection using the Qiagen EndoFree Plasmid Maxi kit. Each purified vector was used for three transfections and purification from glycerol stock was repeated, as needed.

#### Transfection of primary astrocytes

Primary astrocytes were seeded in three 24-well plates at varying densities so that they would reach 70-80% confluence on three different days for independent transfections. Once each plate reached 70-80% confluence, the cells were transfected in triplicate using optimized conditions to deliver 750 ng pGL4.10[*luc2*] firefly luciferase reporter vector DNA (unmodified, modified with promoter only, or modified with putative enhancer region and promoter) and 75 ng pRL-TK renilla luciferase reporter vector DNA. Three wells were also treated with only Opti-MEM and transfection reagents to serve as a mock transfected control. The cells were then cultured for approximately 22 hours in a humidified incubator at 37°C with 5% CO_2_. This transfection process was repeated two more times with freshly thawed primary astrocytes with matched passage numbers and freshly purified vectors so that 9 independent transfections were completed.

#### Transfection of HEK293Ts

HEK293Ts were seeded in three 24-well plates at varying densities so that they would reach 70-90% confluence on three different days for independent transfections. Once each plate reached 70-90% confluence, the cells were transfected in triplicate with 500 ng pGL4.10[*luc2*] firefly luciferase reporter vector DNA (unmodified, modified with promoter only, or modified with putative enhancer region and promoter) and 50 ng pRL-TK renilla luciferase reporter vector DNA with 1 µL P3000 Reagent and 0.75 µL Lipofectamine 3000 diluted in Opti-MEM for a total volume of 50 µL/well. Three wells were also treated with only Opti-MEM and transfection reagents to serve as a mock transfected control. The cells were then cultured for approximately 24 hours in a humidified incubator at 37°C with 5% CO_2_. This transfection process was repeated two more times with freshly thawed HEK293Ts with matched passage numbers and freshly purified vectors so that 9 independent transfections were completed.

#### Luciferase assay

Luciferase assay reagents were prepared using the Promega Dual-Luciferase Reporter Assay System, according to manufacturer’s instructions. After transfection with luciferase reporter vectors, primary astrocytes were washed with PBS, incubated in 500 µL Passive Lysis Buffer/well with rocking at room temperature for 15 minutes, and then gently pipetted to aid lysis with mechanical force. After transfection with luciferase reporter vectors, HEK293Ts were washed with PBS and lysed in 500 µL Passive Lysis Buffer/well with rocking at room temperature for 10 minutes. Each lysate was then collected and vortexed for 10 seconds. 20 µL of each lysate was added to a white, flat-bottom 96-well plate in triplicate for a total of 9 wells/condition. 20 µL Passive Lysis Buffer was also added to 9 wells to serve as a negative control. Each well was assayed using a SpectraMax iD5 Multi-Mode Microplate Reader by injecting 100 µL Luciferase Assay Reagent II, waiting 2 seconds, measuring firefly luciferase fluorescence for 10 seconds, injecting 100 µL Stop & Glo Reagent, waiting 2 seconds, and measuring renilla luciferase fluorescence for 10 seconds.

#### Generation of rs7132908 risk allele ESC clones

A guide RNA and homology-directed repair template were designed to change the rs7132908 non-risk G allele to the obesity risk A allele with CRISPR-Cas9 in the ESC model. These methods were adapted from a previously published protocol for highly efficient CRISPR-Cas9 editing in human stem cells^78^. The guide RNA was designed with the help of the CRISPOR program (RRID: SCR_015935)^79^. The guide RNA was prepared by incorporating the 20 bp target sequence into two 60-mer oligos purchased as 25 nmole DNA oligos from IDT which were then annealed, amplified with PCR using Phusion High-Fidelity DNA polymerase, and purified with extraction with the Takara NucleoSpin Gel and PCR Clean-Up kit after visualization with gel electrophoresis. The guide RNA was then cloned into the gRNA_Cloning vector^80^, which was a gift from George Church (Addgene Cat# 41824; http://n2t.net/addgene:41824; RRID: Addgene_41824), at the AflII restriction site with the NEB Gibson Assembly kit to generate the gRNA_Cloning-rs7132908gRNA vector. The homology-directed repair template was prepared by designing a 100 bp single-stranded oligonucleotide centered around the gRNA sequence and with the desired base change, which was then purchased as a 4 nmole Ultramer DNA oligo from IDT. 0.5 µg gRNA_Cloning-rs7132908gRNA vector, 0.5 µg pCas9_GFP vector^81^, which was a gift from Kiran Musunuru (Addgene Cat# 44719; http://n2t.net/addgene:44719; RRID: Addgene_44719), and 1 µg homology-directed repair template/well were transfected into 70-80% confluent ESCs on irradiated MEFs in a 6-well plate with 3 µL/well Lipofectamine Stem in 50 µL DMEM/F12. The cells were cultured in a humidified incubator at 37°C with 5% CO_2_ for 48 hours. After transfection, single cells were lifted and 5,000-15,000 GFP+ cells were sorted into a 10 cm dish coated with Matrigel Growth Factor Reduced Basement Membrane Matrix diluted in IMDM and MEFs with fluorescence-activated cell sorting. After 10-15 days of maintenance, individual clones were manually picked and used for both screening and expansion. Some cells from each clone were used for Proteinase K DNA extraction. This DNA was used as a template for PCR across the edited region using the Phusion High-Fidelity DNA polymerase and the PCR products were then used for both restriction digestion screening and Sanger sequencing to confirm the base change (**Fig. S5A-B**). Restriction digestion was a possible screening method because the change from the rs7132908 non-risk G allele to obesity risk A allele generated a unique BfaI restriction site. Electropherograms and sequence files produced from Sanger sequencing were analyzed using SnapGene software (RRID: SCR_015052). Clones confirmed to be homozygous for the rs7132908 obesity risk A allele underwent further validation with karyotyping (**Fig. S5C**), *de novo* CNV analysis (**Supp. Table 11**), mycoplasma contamination testing (**Fig. S5D**), and Sanger sequencing at the top 10 most likely off-target sites (**Supp. Table 12**).

#### Karyotyping

ESCs were passaged into a T25 flask and cultured under normal conditions until the cells reached 60-70% confluence. The flask was then packaged and shipped to Cell Line Genetics for G-band karyotyping of live cultures. Karyotyping reports indicated that all ESC lines had a normal human female karyotype (**Fig. S5C**).

#### DNA extraction from cells

To extract DNA from cultured cells for genotyping, PCR, or Sanger sequencing, cells were lifted and then DNA was extracted with the Zymo Quick-DNA Miniprep Plus kit, following manufacturer’s instructions.

#### SNP genotyping

Genome-wide genotyping of DNA from ESC lines for *de novo* CNV analysis was performed using the Illumina Infinium Global Screening Array v3.0 BeadChip genotyping array. Genome-wide genotyping of DNA from human pediatric hypothalamus tissue was performed using the Illumina OmniExpressExome v1.6 BeadChip genotyping array. Genotyping arrays consist of many thousands of short invariant 50mer oligonucleotide probes conjugated to silica beads. Sample DNA is hybridized to the probes and a single-base, hybridization-dependent extension reaction is performed at each target SNP. Arrays are loaded onto an iScan System and scanned to extract data. DMAP files enable identification of bead locations on the BeadChip and quantification of the signal associated with each bead. Alternate alleles (herein denoted A and B) are labeled with different fluorophores. Raw fluorescence intensity from the two-color channels is processed into a discrete genotype call (normalized to continuous value 0-1 B-Allele Frequency (BAF)) and the total intensity from both channels (normalized to continuous value with median=0 Log R Ratio (LRR)) at each SNP which are informative for copy number.

#### Screening for CRISPR off-target effects

The CRISPOR program (RRID: SCR_015935)^79^ was used to identify potential off-target sites for the guide RNA designed to change the rs7132908 non-risk G allele to the obesity risk A allele. Each potential off-target site was ranked by Cutting Frequency Determination score which is used to measure guide RNA specificity. Primers were designed to PCR amplify and Sanger sequence the top 10 potential off-target sites. Six potential off-target sites were excluded from screening because primers could not be designed in these regions with a melting temperature between 56-70°C, likely because these regions were too repetitive. Each potential off-target site was amplified using the Phusion High-Fidelity DNA polymerase. Each PCR product was extracted after visualization with gel electrophoresis with the NEB Monarch DNA Gel Extraction kit to ensure that a fragment of correct length was purified. Each purified PCR product was then Sanger sequenced on both strands. Electropherograms and sequence files produced from Sanger sequencing were analyzed using SnapGene software (RRID: SCR_015052). Sequences from each CRISPR clone were compared to sequences from the parent H9 ESC line to determine that there were no off-target effects in all clones at the top 10 most likely off-target sites (**Supp. Table 12**).

#### Preparation of differentiation medium

Differentiation medium was prepared as previously published^7^ with some modifications. This medium is an optimized, serum-free reformulation of B27 which supports high quality neuronal cultures and overcomes quality variability of B27 due to different sources of bovine serum albumin. A 50X differentiation supplement was prepared containing DMEM/F12 with 1 μg/mL corticosterone, 50 μg/mL linoleic acid, 50 μg/mL linolenic acid, 2.35 μg/mL (±)-α-lipoic acid, 0.32 μg/mL progesterone, 5 μg/mL retinyl acetate, 50 μg/mL (±)-α-tocopherol, 50 μg/mL DL-α-tocopherol acetate, 125 mg/mL bovine serum albumin, 27.15 mg/mL sodium bicarbonate, 3.2 mg/mL L-ascorbic acid, 805 μg/mL putrescine dihydrochloride, 750 μg/mL D(+)-galactose, 250 μg/mL holo-transferrin, 125 μg/mL catalase, 100 μg/mL L-carnitine hydrochloride, 50 μg/mL glutathione, 0.7 μg/mL sodium selenite, 50 μg/mL ethanolamine, 0.1 μg/mL triiodo-L-thyronine sodium salt, and 200 μg/mL insulin. Differentiation medium was then prepared containing DMEM/F12 with 1X differentiation supplement, 1X Antibiotic-Antimycotic, 1X GlutaMAX, and 2.5 μg/mL superoxide dismutase.

#### Differentiation to hypothalamic neural progenitors

ESCs were plated as single cells at 1 million cells/well in a matrigel-coated 6-well plate or 200,000 cells/well in a matrigel-coated 24-well plate and cultured in mTeSR1 medium with 10 μM ROCK Inhibitor Y-27632 for 24 hours in a humidified incubator at 37°C with 5% CO_2_. After 24 hours, on day 0, the medium was changed to differentiation medium with 1 μM LDN-193189 and 10 μM SB-431542 for dual SMAD inhibition. On days 2, 4, 6, and 8, the medium was changed to differentiation medium with 1 μM LDN-193189, 10 μM SB-431542, 1 μM SAG, 1 μM Purmorphamine, and 10 μM IWR-1-endo for dual SMAD and Wnt signaling inhibition and Shh activation. This method directed the ESCs toward ventral diencephalon forebrain cell identity. On days 9, 11, and 13, the medium was changed to differentiation medium with 10 μM DAPT and 0.01 μM retinoic acid to direct the cells to exit cell cycle. Hypothalamic neural progenitors were collected for downstream experiments on day 14 (**Fig. 3A**). These methods were previously optimized and validated^7^. To confirm hypothalamic neural progenitor identity, we performed immunohistochemistry and observed expected expression of NKX2-1, which is a marker for the developing hypothalamus^82^ (**Fig. S1B**), and NeuN, which is a marker for post-mitotic neurons (**Fig. S1C**).

#### Differentiation to hypothalamic neurons

On day 14, hypothalamic neural progenitors were washed with DPBS, incubated at 37°C with Accutase for up to 7 minutes, treated with 2 volumes of medium to neutralize the Accutase, pelleted by centrifugation at 200 rcf for 3 minutes at 25°C, resuspended in differentiation medium with 10 ng/mL BDNF, and seeded at 1 million cells/well in a laminin-coated 6-well plate or 200,000 cells/well in a laminin-coated 24-well plate. Laminin-coated plates were prepared by diluting laminin to 0.05 mg/mL in cold Hanks’ Balanced Salt Solution, distributing 10 mL laminin solution across each plate, incubating overnight at 4°C, incubating at 37°C for 2 hours before use, and washing with PBS 3 times before use. The medium was replaced with fresh differentiation medium with 10 ng/mL BDNF every 2-3 days until day 40 to promote hypothalamic neuron maturation. These methods were previously optimized and validated^7^.

#### Fluorescent immunohistochemistry

Cells for immunohistochemistry were cultured on acid-treated #1.5 glass coverslips. The cells were washed with PBS, fixed with 4% paraformaldehyde for 10 minutes at room temperature, and then incubated with PBS for 5 minutes at room temperature three times to wash. The cells were incubated in blocking solution (PBS with 5% (w/v) bovine serum albumin and 0.3% Triton X-100) for 1 hour at room temperature. After blocking, primary antibodies (Anti-MAP2 (Abcam Cat# ab5392; RRID: AB_2138153), Anti-NKX2-1 (Cell Marque Cat# 343M-95; RRID: AB_1158934), and Anti-NeuN (Millipore Sigma Cat# MAB377; RRID: AB_2298772)) diluted in blocking solution (1:500) were added to the cells, then incubated overnight at 4°C with gentle rocking. After the primary antibody incubation, the cells were incubated with PBST for 10 minutes at room temperature three times to wash. Appropriate secondary antibodies (Anti-Chicken (Abcam Cat# ab150169; RRID: AB_2636803) and Anti-Mouse (Invitrogen Cat# A-11001; RRID: AB_2534069)) diluted in blocking solution (1:500) were added to the cells, then incubated for 1 hour at room temperature, protected from light. After the secondary antibody incubation, the cells were incubated with PBST for 5 minutes at room temperature three times to wash. The cells were then washed with PBS for 3 minutes at room temperature and incubated with 300 nM DAPI for 5 minutes at room temperature to stain nuclei. After DAPI incubation, the cells were washed with PBS three times. The glass coverslips were mounted on glass slides with ProLong Gold Antifade Mountant. The cells were visualized with an Olympus DP74 camera using appropriate fluorescent filters and Olympus cellSens Standard software. Images for each fluorescent channel were merged using ImageJ (RRID: SCR_003070)^83^.

#### Nuclei isolation

After hypothalamic neuron differentiation, the cells were washed with PBS, incubated at 37°C with Accutase for up to 7 minutes, treated with 2 volumes of medium to neutralize the Accutase, and pelleted by centrifugation at 300 rcf for 5 minutes at 4°C. The cell pellet was resuspended in PBS with 0.04% bovine serum albumin. 1 million cells or less were pelleted by centrifugation at 300 rcf for 5 minutes at 4°C and then resuspended in 100 μL chilled lysis buffer (water with 10 mM Trizma hydrochloride, 10 mM sodium chloride, 3 mM magnesium chloride, 1% bovine serum albumin, 0.1% Tween-20, 1 mM DTT, 1 U/μL RNase inhibitor, and 0.1% IGEPAL CA-630). The cells were incubated in lysis buffer on ice for 1 minute and then 500 μL chilled wash buffer (water with 10 mM Trizma hydrochloride, 10 mM sodium chloride, 3 mM magnesium chloride, 1% bovine serum albumin, 0.1% Tween-20, 1 mM DTT, and 1 U/μL RNase inhibitor) was added. The nuclei were pelleted by centrifugation at 500 rcf for 5 minutes at 4°C. Addition of chilled wash buffer and pelleting were repeated two more times. The nuclei were then resuspended in chilled nuclei buffer (water with 1X Nuclei Buffer, 1 mM DTT, and 1 U/μL RNase inhibitor) to a concentration of 8,000 nuclei/μL in at least 25 μL and strained using a 35 μm strainer.

#### Single-nucleus RNA-seq and ATAC-seq library preparation

Single-nucleus RNA-seq and ATAC-seq libraries were prepared using the 10X Genomics Chromium Single Cell Multiome ATAC + Gene Expression workflow. Libraries were prepared from the rs7132908 non-risk G allele cells from two independent differentiations (biological replicates) for a total of 4 technical replicates and from the rs7132908 risk A allele cells from two CRISPR clones (biological replicates) and three independent differentiations (biological replicates) for a total of 4 technical replicates. In brief, isolated nuclei in chilled nuclei buffer were transposed in bulk which simultaneously fragmented DNA in regions of open chromatin and added adapter sequences to the ends of the DNA fragments. The transposed nuclei were then loaded onto a microfluidic chip which was run in the Chromium Controller instrument. In the instrument, nuclei were individually partitioned with Gel Beads-in-emulsion (GEMs). Each Gel Bead contains oligonucleotides with a unique 16 bp 10X Barcode sequence, a poly(dT) sequenced to capture mRNA, and a Spacer sequence that enables barcode attachment to transposed DNA fragments. The GEMs were then incubated to attach unique 10X Barcodes to mRNA and transposed DNA fragments which served to associate mRNA and transposed DNA fragments back to the same nucleus. Unique molecular identifiers (UMIs) were also used to distinguish individual, captured mRNA molecules for quantification. A reverse transcription reaction converted the mRNA into full-length cDNA. The GEMs were then broken and pooled fractions were recovered and purified. The products were taken through a pre-amplification PCR step to fill gaps and ensure maximum recovery of barcoded ATAC and cDNA fragments. The pre-amplified products were then used as input for both ATAC-seq library preparation and cDNA amplification for RNA-seq library preparation. Completed RNA-seq libraries were quantified and assessed with Agilent High Sensitivity D1000 ScreenTape assays and ATAC-seq libraries were quantified and assessed with Agilent High Sensitivity D5000 ScreenTape assays. RNA-seq libraries were then pooled and sequenced on the Illumina NovaSeq 6000 platform to reach a minimum of 20,000 paired-end reads/nucleus. ATAC-seq libraries were then pooled and sequenced on the Illumina NovaSeq 6000 platform to reach a minimum of 25,000 paired-end reads/nucleus.

#### cDNA generation

RNA samples were quantified with Qubit RNA High Sensitivity Assays. 30 ng of each RNA sample was used for cDNA generation using SuperScript IV VILO Master Mix after treatment with ezDNase to remove any DNA contamination. No reverse transcriptase controls were also generated using SuperScript IV VILO ‘No RT’ Control Master Mix.

#### Quantitative real-time polymerase chain reaction (RT-qPCR)

TaqMan Gene Expression Assays for *FAIM2* and human *18S* ribosomal RNA were validated with standard curves generated by pooling all cDNA samples quantified in an experiment to represent average conditions of all samples. The *FAIM2* standard curve consisted of 5 points generated by a 1:5 serial dilution ranging from 0.0024 to 1.5 ng in triplicate. The *18S* standard curve consisted of 8 points generated by a 1:5 serial dilution ranging from 0.0000192 to 1.5 ng in triplicate. Each sample was quantified with TaqMan Fast Advanced Master Mix and the Agilent AriaMX Real-Time PCR System. After assay validation, 0.5 ng of each experimental cDNA sample and no reverse transcriptase control were assayed in duplicate. Additionally, no template controls were assayed in triplicate.

### Quantification and statistical analysis

#### Prediction of risk allele’s effect on transcription factor binding

The genomic position and alternative allele of rs7132908 (determined using SNPlocs.Hsapiens.dbSNP155.GRCh38 and BSgenome R packages) were used to scan through all position frequency matrix databases using the R package MotifDb to identify potential transcription factor binding disruption effects. The motifbreakR function^84^ was used with parameters filter=TRUE, threshold=0.0005, method=’ic’, bkg=c(A=0.25, C=0.25, G=0.25, T=0.25), and BPPARAM=BiocParallel::SerialParam().

#### GWAS-eQTL colocalization

Childhood obesity GWAS summary statistics from the European ancestry population in the EGG consortium were used. Common variants (minor allele frequency ≥ 0.01) from the 1000 Genomes Project (v3)^85^ were used as a reference panel. SNP-gene sets from our variant-to-gene mapping efforts were used as leads. We used ColoQuiaL^86^ to test genome-wide colocalization of each lead against GTEx eQTLs (v8) (RRID: SCR_013042)^36^ from all 49 available tissues. Evidence of colocalization between a given childhood obesity GWAS signal and eQTL signal was identified by a conditional posterior probability of colocalization ≥ 0.8.

#### Luciferase assay data analysis

All fluorescence values were reduced by the average signal in the 9 negative control wells to correct for background fluorescence in the Passive Lysis Buffer and 96-well plate. The firefly luciferase fluorescence signal was then divided by the renilla luciferase fluorescence signal in each well to adjust for sample-to-sample variability due to differences in cell numbers, transfection efficiency, and pipetting. Normalized firefly luciferase fluorescence values were averaged for each condition (n=9). Normalized fold change was calculated by dividing the average normalized firefly luciferase fluorescence values for each condition by this value produced by the promoter only vector (pGL4.10[*luc2*]-*FAIM2,* pGL4.10[*luc2*]-*LIMA1,* or pGL4.10[*luc2*]-*RACGAP1*).

Assays were excluded from statistical analysis if there was fluorescence detected (normalized fold change > 0.1) in the negative control condition or if at least one normalized fold change value was greater than 2 standard deviations away from the mean of all other assays performed. Multiple independent transfections and assays were performed and are stated in the figure legend. All data are represented as mean ± standard deviation. Statistical analyses and visualization were performed using GraphPad Prism (RRID: SCR_002798) and ordinary one-way ANOVA tests with Tukey’s correction for multiple comparisons. *P*-values < 0.05 were considered significant. \**P*-value < 0.05, \*\**P*-value < 0.01, \*\*\**P*-value < 0.001.

#### CNV detection

Samples must meet minimum quality control standards of call rate > 98% and LRR standard deviation < 0.3 to be used for CNV detection. We used PennCNV (RRID: SCR_002518) as our main CNV detection algorithm of the Illumina Infinium Global Screening Array v3.0 data due to its widespread usage. We filtered PennCNV calls to include CNVs with number of SNPs supporting ≥20, length ≥100,000, and Segmental Duplication track coverage < 0.5. Related cell line clone CNV calls were compared to ensure consistency in CNV calling. All genomic coordinates are in human genome build version GRCh37.

#### *De novo* CNV detection

The related cell line clones annotated for each sample were verified by pairwise comparison of genome-wide SNP genotyping content using PLINK (RRID: SCR_001757). The “child” cell line CNVs were compared to their corresponding “parent” cell line CNVs using bedtools and if at least 50% reciprocal overlap is not observed, annotated as *de novo*. Such putative *de novo* calls were BAF LRR plotted for each pair of “child” and “parent” to allow for side-by-side comparison to ensure the *de novo* was not an erroneous call.

#### Bulk RNA-seq analysis

Sequencing data was demultiplexed to generate FASTQ files using Illumina bcl2fastq2 Conversion Software (RRID: SCR_015058). FASTQ files were assessed with FastQC (RRID: SCR_014583)^87,88^ to verify that there was high sequence quality, expected sequence length, and no adapter contamination. Paired-end FASTQ files for each replicate of primary astrocytes were mapped to the human reference genome (GRCh38) using STAR (RRID: SCR_004463)^89^. Genes were annotated using GENCODE human release 40 (RRID: SCR_014966)^90^. Raw read counts were calculated using HTSeq-count (RRID: SCR_011867)^91^. Paired-end FASTQ files for each replicate of all other cell types and tissue were mapped to the Ensembl human reference transcriptome (GRCh38)^92^ using Kallisto (RRID: SCR_016582)^93,94^. Abundance data generated with Kallisto was read into R (RRID: SCR_001905) using the package tximport (RRID: SCR_016752)^95^, annotated with Ensembl human gene annotation data (version 86)^92^ using ensembldb (RRID: SCR_019103)^96^ and EnsDb.Hsapiens.v86, and summarized as counts per million (cpm) at the gene level using edgeR (RRID: SCR_012802)^97^. Genes with less than 1 cpm in 2 or 3 samples, depending on the smallest set of replicates in the analysis, were removed to increase statistical power to detect differentially expressed genes. Samples within each analysis were normalized with the trimmed mean of M values (TMM) method^98^. The R package limma (RRID: SCR_010943)^99^ was used to identify differentially expressed genes by first applying precision weights to each gene based on its mean-variance relationship using the voom function and then linear modeling and Bayesian statistics were employed to detect genes that were up- or down-regulated in each condition. Genes with an adjusted *P*-value < 0.05 and |log2 fold change| > 0.58 were considered significantly differentially expressed. Coordinates for the rs7132908 TAD were determined using the TADKB database^51^ and considering the most conservative region documented in all reported human cell lines (GRCh37). A list of genes in the rs7132908 TAD region were exporting using the UCSC Genome Browser (GRCh37) (RRID: SCR_005780)^100,101^. Significantly differentially expressed genes were clustered using Pearson correlation and the R function hclust. The clustered genes were cut into 2 modules in ESCs and 5 modules in hypothalamic neural progenitors. Significantly enriched Gene Ontology terms^52,53^ in each module were identified using gprofiler2 (RRID: SCR_018190)^102,103^. Results were visualized using ggplot2 (RRID: SCR_014601)^104^, gplots, and plotly.

#### Bulk ATAC-seq analysis

Sequencing data was demultiplexed to generate FASTQ files using Illumina bcl2fastq2 Conversion Software (RRID: SCR_015058). ATAC-seq peaks were called following the ENCODE ATAC-seq pipeline (https://www.encodeproject.org/atac-seq/). Briefly, paired-end reads from three replicates for each cell type were aligned to the human reference genome (GRCh38) using bowtie2 (RRID: SCR_016368)^105^, and duplicate reads were removed from the alignment using Picard (RRID: SCR_006525) MarkDuplicates and SAMtools (RRID: SCR_002105)^106^. Narrow peaks were called independently for each replicate using MACS2^107^ with parameters -p 0.01 --nomodel --shift -75 --extsize 150 -B --SPMR --keep-dup all --call-summits. Reproducible peaks, peaks called in at least 2 replicates (with at least 1 bp overlap), were used to generate a consensus set of peaks. Signal peaks were normalized using csaw^108^ in 10 kilobase (kb) bin background regions. A threshold of cpm > 1 was used to exclude peaks with low abundance from the analysis. Tests for differential accessibility between rs7132908 genotypes were conducted with the glmQLFit approach implemented in edgeR (RRID: SCR_012802)^97^ using the normalization factors calculated by csaw. Open chromatin regions with adjusted *P*-value < 0.05 and |log2 fold change| > 1 were considered differentially accessible. Results were visualized using ggplot2 (RRID: SCR_014601)^104^.

#### Hi-C analysis

Hi-C analysis was performed as previously described^47^. In brief, sequencing data was demultiplexed to generate FASTQ files using Illumina bcl2fastq2 Conversion Software (RRID: SCR_015058). Paired-end reads from each replicate were pre-processed using the HiCUP pipeline (RRID: SCR_005569)^109^ and aligned to the human reference genome (GRCh38) with bowtie2 (RRID: SCR_016368)^105^. The alignments files were parsed to pairtools (RRID: SCR_023038)^110^ to process and pairix^111^ to index and compress, then converted to Hi-C matrix binary format (.cool) by cooler^112^ at multiple resolutions (500 bp, 1, 2, 4, 10, 40, 500 kb and 1 megabase (Mb)) and normalized with the ICE method^113^. The matrices from different replicates were merged at each resolution using cooler^112^. Mustache^114^ and Fit-Hi-C2^115^ were used to call significant intra-chromosomal interaction loops from merged replicates matrices at three resolutions (1 kb, 2 kb, and 4 kb), with significance thresholds of q-value < 0.1 and *P*-value < 1×10^−6^. The identified interaction loops were merged between both tools at each resolution. Lastly, interaction loops from all three resolutions were merged with preference for smaller resolution if there was overlap.

#### Single-nucleus RNA-seq and ATAC-seq pre-processing

Cell Ranger ARC analysis pipelines were used to process sequencing data generated with the 10X Genomics Chromium Single Cell Multiome ATAC + Gene Expression workflow. Sequencing data was demultiplexed to generate FASTQ files using mkfastq. The FASTQ files were aligned to the GRCh38 human reference genome with the Cell Ranger ARC package (RRID: SCR_023897) and cells were called using parameters -count --min-atac-count=2000 --min-gex-count=1000.

66,120 cells homozygous for the rs7132908 non-risk G allele representing two separate differentiations were sequenced. 45,916 cells homozygous for the rs7132908 obesity risk A allele representing two different clonal lines and three different differentiations were also sequenced. All 112,036 cells then underwent quality control to remove ambient RNA using SoupX (RRID: SCR_019193)^116^ with the contamination fraction automatically estimated for each sample and the count matrices were re-adjusted after removal. Doublets were detected and removed using the Python package Scrublet (RRID: SCR_018098)^117^, and cells with >10% mitochondrial reads were filtered out using Seurat (RRID: SCR_016341)^118^. After quality control, we retained 71,818 cells for downstream analyses.

RNA-seq data from all samples was SCTransformed (RRID: SCR_022146)^119,120^, integrated using the IntegrateData function, and then batch corrected using Harmony (RRID: SCR_022206)^121^ for differentiation, biological, and technical replicates. PCA and UMAP reduction were performed using the first 30 empirically selected principal components with standard pipelines (**Fig. S2A-C**).

We ran peak calling using MACS3 (https://macs3-project.github.io/MACS/) for each sample with their corresponding ATAC-seq fragments files. Peaks from all samples were pooled and reduced to a final set of 383,029 peaks accessible in at least one sample. This peak set was used to create a ChromatinAssay using Signac (RRID: SCR_021158)^122^. The peaks were filtered through ENCODE hg38 blacklist regions (https://github.com/Boyle-Lab/Blacklist/blob/master/lists/hg38-blacklist.v2.bed.gz) and annotated with EnsDb.Hsapiens.v86. We performed quality control following metrics recommended by Signac^122^, including nucleosome banding pattern, TSS enrichment score, total number of fragments in peaks, fraction of fragments in peaks, and ratio of reads in genomic blacklist regions; we removed cells that were outliers by these metrics. We performed term frequency-inverse document frequency normalization with the RunTFIDF function and feature selection and dimension reduction using singular value decomposition (SVD) on the TD-IDF matrix with the RunSVD function, which produced latent semantic indexing components (LSI)^123^. Uniform manifold approximation and projection embedding was computed based on the first 29 LSI components (second to the 30^th^) for visualization in two-dimensional space with the RunUMAP function. The first component, being in strong correlation with total counts, was not used. Results were visualized using Seurat^118^ and ggplot2 (RRID: SCR_014601)^104^.

#### Single-nucleus RNA-seq cell type identification

A previously published human hypothalamic arcuate nucleus single-cell RNA-seq dataset^54^ was used as a reference dataset to identify cell types in our single-nucleus RNA-seq dataset. Pairwise correspondences or ‘anchors’ between individual cells in each dataset were defined using the Seurat (RRID: SCR_016341) function FindTransferAnchors^55^. Then each cell in our dataset was classified as one of the cell types in the reference dataset (neuron, astrocyte, OPC, mature oligodendrocyte, microglia, ependymal, pericyte, immature oligodendrocyte, fibroblast, choroid, and tanycyte) using the Seurat function TransferData^55^, where the reference cell type with the highest observed classification score was assigned. As a result, neuron, astrocyte, OPC, ependymal, fibroblast, and tanycyte annotations were added to our dataset (**Fig. S2D**). We then prioritized cells with a classification score ≥ 0.8 for downstream analyses as this threshold has been previously demonstrated to increase accuracy^55^. In summary, we identified 38,044 cells as neurons, OPCs, or fibroblasts with a classification score above our threshold. PCA and UMAP reduction were performed using the first 20 empirically selected principal components with standard pipelines (**Fig. 4A**). All cells annotated as neurons were then subset and reclustered with PCA and UMAP reduction using the first 15 empirically selected principal components (**Fig. 4C**). Results were visualized using Seurat^118^ and ggplot2 (RRID: SCR_014601)^104^.

#### Transcriptome correlation with pediatric hypothalamus tissue and GTEx RNA-seq data

Pseudobulk TPMs were calculated for each annotated cell type and replicate sample in the single-nucleus RNA-seq dataset by normalizing SoupX-corrected counts by gene size using gene annotation data from GENCODE human release 38 (GRCh37) (RRID: SCR_014966)^90^ and previously published code^124^. TPMs from all rs7132908 non-risk allele replicate samples for each annotated cell type were then averaged. Similarly, average TPMs were also calculated for the rs7132908 non-risk allele replicate samples in the bulk RNA-seq datasets generated from the hypothalamic neural progenitors and human pediatric hypothalamus tissue sequenced inhouse. Median gene-level TPM data by tissue was downloaded from the GTEx Analysis RNA-seq database (v8) (RRID: SCR_013042)^36^. Ensembl gene IDs with version suffixes were converted to gene names using gene annotation data from GENCODE human release 26 (GRCh37) (RRID: SCR_014966)^90^. Average TPMs for each cell type of interest were merged with average TPMs from the human pediatric hypothalamus tissue and GTEx data. Then, the spearman rank correlation of genes expressed at greater than 5 TPMs in at least 2 samples were calculated using the R (RRID: SCR_001905) cor function. *P*-values for each correlation were calculated using the R cor.test function. Results were visualized in dot plots using ggplot2 (RRID: SCR_014601)^104^.

#### Neuron transcriptome comparison to human prenatal hypothalamus tissue

To compare the transcriptome of the cells annotated as neurons in the single-nucleus RNA-seq dataset to human prenatal hypothalamic nuclei, data from the Allen Brain Atlas^58–61^ was downloaded as upregulated gene sets from the Harmonizome database^125^. Left and right hemisphere gene sets for each hypothalamic nucleus were combined and used for downstream analysis. To infer the average expression of each gene set per single cell in the neuron dataset compared to random control genes, module scores for each gene set were calculated using the Seurat (RRID: SCR_016341) function AddModuleScore^126^. Average module scores per neuron cluster were plotted as the column Z-score for visualization. Results were visualized using ggplot2 (RRID: SCR_014601)^104^.

#### Single-nucleus RNA-seq differential expression analyses

Differential expression analysis of single-nucleus RNA-seq data was performed with DESeq2 (RRID: SCR_015687)^127^, following the standard workflow. In brief, raw counts and appropriate metadata for cell aggregation and comparison were extracted and used to create a SingleCellExperiment object using the R package SingleCellExperiment^128,129^. Counts were aggregated to the sample level for each cell type using the Matrix.utils function aggregate.Matrix. DESeq2 objects were created from the raw counts, appropriate metadata, and design formula to compare the rs7132908 obesity risk allele to the non-risk allele in each cell type using the DESeq2 function DESeqDataSetFromMatrix^127^. Differential expression analysis in each cell type was run using the DESeq2 function results^127^ and an adjusted *P*-value threshold of 0.05. The resulting log2 fold changes were shrunk using the apeglm method^130^. Genes with an adjusted *P*-value < 0.05 and |log2 fold change| > 0.58 were considered significantly differentially expressed. Results were visualized in volcano plots using ggplot2 (RRID: SCR_014601)^104^. Significantly differentially expressed genes were clustered using the R (RRID: SCR_001905) function hclust and plotted in heatmaps using the R package pheatmap (RRID: SCR_016418). Significantly enriched Gene Ontology terms^52,53^ in each set of genes significantly up- or down-regulated in each cell type were identified using gprofiler2 (RRID: SCR_018190)^102,103^.

#### Single-nucleus ATAC-seq differential accessibility analyses

To find differentially accessible regions due to rs7132908 genotype, we performed differential accessibility tests between cells homozygous for either rs7132908 allele. We implemented logistic regression using the FindMarkers function from Signac (RRID: SCR_021158)^122^, with the total number of fragments in peaks as a latent variable to mitigate the effect of differential sequencing depth and using a min.pct threshold of 0.01 due to sparse single-nucleus ATAC-seq data. To ensure data correspondence, we used only the 38,044 annotated cells that had a classification score ≥ 0.8 by the RNA-seq analysis for this differential accessibility analysis. *P*-value adjustment was performed internally using Bonferroni correction based on the total number of peaks in the dataset. We repeated this analysis for each annotated cell type: neurons, OPCs, and fibroblasts.

We performed DNA motif analysis to identify potentially important genotype-specific regulatory sequences in different groups of differentially accessible peaks. We used motif position frequency matrices from the JASPAR 2022 CORE collection database^131^. The FindMotifs function from Signac^122^ performed hypergeometric test on these differentially accessible peaks to test the probability of observing the motif at the given frequency by chance, compared to a background set of peaks matched for GC content.

#### Quantitative real-time polymerase chain reaction (RT-qPCR) analysis

Cq values for each sample were determined with the Agilent Aria software. To validate each TaqMan Gene Expression Assay using a standard curve, Cq values from each triplicate of samples were averaged and then plotted against the log of their corresponding mass of cDNA input (ng) using Microsoft Excel (RRID: SCR_016137). A linear trendline was then added to each graph and the R^2^ values and linear equations were displayed. Primer efficiency was calculated with 10^(-1/slope)^. Percent primer efficiency was calculated by dividing the primer efficiency by 2. TaqMan Gene Expression Assays passed standard curve validation if the R^2^ value was greater than 0.99 and the percent primer efficiency was between 90-110%. Assays were used to calculate normalized relative expression if the no reverse transcriptase and no template control samples did not generate a Cq value. Normalized relative expression was calculated using (E*_FAIM2_*^((mean *FAIM2* Cq in non-risk cells on day 0) – (mean *FAIM2* Cq in experimental sample))^)/(E*_18S_*^((mean *18S* Cq in non-risk cells on day 0) – (mean *18S* Cq in experimental sample))^), where E is primer efficiency. Results were visualized using GraphPad Prism (RRID: SCR_002798). Independent differentiations were performed and are represented by individual points on each graph. All data are represented as mean ± standard deviation.

